# Genome-wide patterns of copy number variation in the diversified chicken genomes using next-generation sequencing

**DOI:** 10.1101/002006

**Authors:** Guoqiang Yi, Lujiang Qu, Jianfeng Liu, Yiyuan Yan, Guiyun Xu, Ning Yang

**Affiliations:** Department of Animal Genetics and Breeding, College of Animal Science and Technology, China Agricultural University, Beijing, China

**Keywords:** Copy number variation, Whole genome sequencing, aCGH, Genetic diversity, Chicken

## Abstract

Copy number variation (CNV) is important and widespread in the genome, and is a major cause of disease and phenotypic diversity. Herein, we perform genome-wide CNV analysis in 12 diversified chicken genomes based on whole genome sequencing. A total of 9,025 CNV regions (CNVRs) covering 100.1 Mb and representing 9.6% of the chicken genome are identified, ranging in size from 1.1 to 268.8 kb with an average of 11.1 kb. Sequencing-based predictions are confirmed at high validation rate by two independent approaches, including array comparative genomic hybridization (aCGH) and quantitative PCR (qPCR). The Pearson’s correlation values between sequencing and aCGH results range from 0.395 to 0.740, and qPCR experiments reveal a positive validation rate of 91.71% and a false negative rate of 22.43%. In total, 2,188 predicted CNVRs (24.2%) span 2,182 RefSeq genes (36.8%) associated with specific biological functions. Besides two previously accepted copy number variable genes *EDN3* and *PRLR*, we also find some promising genes with potential in phenotypic variants. *FZD6* and *LIMS1*, two genes related to diseases susceptibility and resistance are covered by CNVRs. Highly duplicated *SOCS2* may lead to higher bone mineral density. Entire or partial duplication of some genes like *POPDC3* and *LBFABP* may have great economic importance in poultry breeding. Our results based on extensive genetic diversity provide the first individualized chicken CNV map and genome-wide gene copy number estimates and warrant future CNV association studies for important traits of chickens.

## Introduction

Copy number variation (CNV) is defined as gains or deletions of DNA fragments of 50 bp or longer in length in comparison with reference genome (Redon et al. 2006; Bickhart et al. 2012). CNVs contribute significantly to both disease susceptibility and resistance and normal phenotypic variability in humans (McCarroll and Altshuler 2007; Zhang et al. 2009; Altshuler et al. 2010) and animals (Liu et al. 2010; Yalcin et al. 2011; Wang et al. 2012a; Wang et al. 2012b). Four major mechanisms have been found to be related to CNV formation including non-allelic homologous recombination (NAHR), non-homologous end joining (NHEJ), Fork Stalling and Template Switching (FoSTeS) and LINE1 Retrotransposition (Hastings et al. 2009; Zhang et al. 2009). Additionally, segmental duplications (SDs) which are duplicated sequences (insertions) of ≥ 1 kb in length and ≥ 90% sequence identity are also suggested to be a major catalyst and hotspot for CNV formation (Sharp et al. 2005; Alkan et al. 2009), mainly because regions flanking by SDs are susceptible to recurrent rearrangement by NAHR (Sharp et al. 2005; Freeman et al. 2006). In terms of total bases involved, the percentage of the genome affected by CNVs is higher than that of single nucleotide polymorphism (SNP) markers. Although SNPs are generally considered as suitable markers in the genome-wide association studies (GWAS), most reported SNP variants have relatively limited effects and explain only a small proportion of phenotypic variance (Manolio et al. 2009). Further, CNVs encompassing part or all of a gene or regulatory elements are believed to have potentially larger effects by influencing gene expression indirectly through changing gene structure and dosage, altering gene regulation, exposing recessive alleles and other mechanisms (Redon et al. 2006; Zhang et al. 2009; Conrad et al. 2010; Liu and Bickhart 2012). CNVs are also found to alone capture 18% to 30% of the total detected genetic variation in gene expression in humans and animals, and might contribute to a fraction of the missing heritability (Stranger et al. 2007; Henrichsen et al. 2009). Therefore, identification of CNVs is essential in whole genome fine-mapping of CNVs and association studies for important phenotypes.

Originally, two cost-effective and high-throughput methods including array comparative genomic hybridization (aCGH) and commercial SNP microarrays are used for CNV screening (LaFramboise 2009; Pinto et al. 2011). However, different analytic platforms and tools reveal inconsonant results with minimal overlap owing to different designs and genome coverage or density of probes (Henrichsen et al. 2009; Pinto et al. 2011). Due to the limitation in resolution and sensibility, it is difficult for the two approaches to detect small CNVs shorter than 1 kb in length and identify the precise breakpoints of CNVs (Bentley et al. 2008; Yoon et al. 2009). Furthermore, the presence of SD regions is a common challenge for the two platforms, because they are often affected by low probe density and cross-hybridization of repetitive sequence (Campbell et al. 2011; Bickhart et al. 2012). Recently, a variety of CNV detection approaches based on next-generation sequencing (NGS) are proposed and offer a promising alternative as they have a higher effective resolution to discover more types and sizes of CNVs (Teo et al. 2012). One effective method is read depth (RD) (also known as depth of coverage (DOC)) with capability of inferring gain or loss of DNA and determining absolute copy number value of each genetic locus, which detects CNVs by analyzing the number of reads that fall in each pre-specified window of a certain size (Abyzov et al. 2011; Szatkiewicz et al. 2013). Hence, the advent of NGS technologies and suitable analytical method promises to systematically identify CNVs at higher resolution and sensitivity.

At present, the three aforementioned high-throughput platforms have been applied to livestock genomics for CNV detection, such as sheep (Norris and Whan 2008), horse (Rosengren Pielberg et al. 2008) and cattle (Bickhart et al. 2012), and suggest several CNVs associated with important phenotypes. CNVs in chickens are also found to be the genetic foundation of phenotypic variation. A duplicated sequence close to the first intron of *SOX5* is associated with the chicken pea-comb phenotype (Wright et al. 2009) and an inverted duplication containing *EDN3* causes dermal hyperpigmentation (Dorshorst et al. 2011). Partial duplication of the *PRLR* also shows to be related to the late feathering (Elferink et al. 2008).

A genome-wide chicken CNV analysis is essential since the chicken is not only an economically important farm animal but also a valuable biomedical model (Wang et al. 2012b; Jia et al. 2013). However, previous CNV studies in chickens based on aCGH and SNP platforms mainly suffered from low resolution and sensitivity (Griffin et al. 2008; Wang et al. 2012b; Crooijmans et al. 2013; Jia et al. 2013; Tian et al. 2013), and a latest report exhibited the detection of four main types of genetic variation from whole genome sequencing data using two chickens (Fan et al. 2013), which suggested the efficiency of CNV detection via deep sequencing. To construct a more refined and individualized chicken CNV map and investigate genome-wide CNV genotyping, benefiting from extensive genetic diversity in Chinese indigenous (Qu et al. 2006) and commercial chickens, we describe the use of NGS data to detect CNVs in the diversified chicken genomes, and estimate genome-wide gene copy number, enabling us to better understand the patterns of CNVs in the chicken genome and future CNV association studies similar to SNPs.

## Results

### Mapping statistics and CNV detection

We performed whole genome sequencing in 12 different breeds of female chickens using Illumina paired-end library and obtained a total of 12.9 Gb of high quality sequence data per individual after quality filtering. After sequence alignment and removing potential PCR duplicates, the sequence depth for each individual varied from 8.2× (CS) to 12.4× (WR), which was sufficient for CNV detection, and the average coverage with respect to the chicken genome reference sequence was 97.2% (**Table 1**). We calculated the average RD for 5 kb non-overlapping windows for all autosomes and performed GC correction as previous reports. The GC-adjusted RD mean and standard deviation (STDEV) of autosomes for each individual was listed in **Table 1**. We applied the program CNVnator to 12 individuals and the average number of CNVs per individual was 1,389, ranging from 703 in WL to 1,975 CNVs in BY. A detailed description of CNV calls could be found in Supplementary Table S1. The mean CNV size in BY (17.4 kb) and CS (14.7 kb) was significantly larger than that of the other individuals (from 4.7 kb in WR to 8.5 kb in SK). In addition, the proportion of CNVs less than 10 kb in length was smaller in BY (52.6%) and CS (54.8%) compared with others (from 73.4% in SK to 90.3% in WR). For all CNVs classified as duplication, the autosomal maximum copy number was 40.8 on chromosome 2 (chr2) in RJF, and the average copy number of all duplicated regions on autosomes in all individuals was 3.88.

**Table 1.** Summary statistics for sequencing and CNVs of individuals

| Chicken abbreviation<sup>a</sup> | Numbers of mapped reads | Depth | Coverage (%) | Autosome reads per 5 kb window<sup>b</sup> | Autosome reads STDEV | Duplications | Deletions | Sequence covered (Mb) |
| --- | --- | --- | --- | --- | --- | --- | --- | --- |
| BY | 102,002,937 | 9.7 | 97.0 | 489.29 | 110.73 | 1,344 | 631 | 34.4 |
| CS | 85,383,494 | 8.2 | 96.9 | 409.93 | 101.42 | 1,166 | 686 | 27.3 |
| DX | 129,847,015 | 12.4 | 97.4 | 623.50 | 130.46 | 592 | 848 | 8.5 |
| LX | 105,152,881 | 10.0 | 97.3 | 503.82 | 112.74 | 935 | 844 | 12.1 |
| RIR | 102,464,756 | 9.8 | 97.3 | 490.96 | 108.21 | 643 | 684 | 8.9 |
| RJF | 105,517,587 | 10.1 | 97.2 | 504.23 | 113.52 | 729 | 641 | 10.1 |
| SG | 85,987,827 | 8.2 | 96.6 | 412.27 | 87.66 | 510 | 568 | 7.5 |
| SK | 95,322,371 | 9.1 | 97.1 | 457.21 | 100.61 | 806 | 692 | 12.7 |
| TB | 107,535,104 | 10.3 | 97.3 | 515.68 | 108.07 | 642 | 705 | 8.8 |
| WC | 119,116,969 | 11.4 | 97.4 | 572.35 | 121.39 | 750 | 803 | 10.2 |
| WL | 118,689,980 | 11.3 | 97.5 | 567.18 | 118.63 | 226 | 477 | 3.7 |
| WR | 130,307,416 | 12.4 | 97.6 | 625.01 | 132.32 | 242 | 508 | 3.5 |

<sup>a</sup>BY, Beijing You; CS, Cornish; DX, Dongxiang; LX, Luxi Game; RIR, Red Island Rhode; RJF, Red Jungle Fowl; SG, Shouguang; SK, Silkie; TB, Tibetan; WC, Wenchang; WL, White Leghorn; WR, White Plymouth Rock.
<sup>b</sup>The number of reads per 5 kb windows after GC correction.

A total of 9,025 CNV regions (CNVRs) allowing for CNV overlaps of 1 bp or greater were obtained, mainly on the 28 autosomes, two linkage groups and sex chromosomes, which amounted to 100.1 Mb of the chicken genome and corresponded to 9.6% of the genome sequence. The individualized chicken CNV map across the genome was shown in **Supplementary Figure S1**. The length of CNVRs ranged from 1.1 to 268.8 kb with an average of 11.1 kb and a median of 6.6 kb. In total, 6,276 (69.5%) out of all CNVRs had size varying from 1.1 to 10 kb (**Figure 1a**). Although chr1 had a maximum of 1,933 CNVRs, the two largest CNVR density, defined as the average distance between CNVRs, were 35.7 kb and 32.0 kb on the chr16 and LGE64 respectively (**Supplementary Table S2**). Meanwhile, Among all CNVRs, 6,160 (68.3%) were present in a single individual, 1,461 (16.2%) were shared in two individuals and 1,404 (15.5%) shared in at least three individuals (**Figure 1b**). Further, the mean and median of the specific CNVRs was 8.9 kb and 5.8 kb in size, whereas the shared CNVRs size was 15.9 kb in average and 9.5 kb as the median. According to the type of CNVRs, they were divided into three categories, including 4,821 gain, 3,854 loss and 350 both (gain and loss) CNVRs. The number of CNVRs in different individuals varied greatly, ranging from 677 in WL to 1,933 in BY, and was positively related to the proportion of specific CNVRs in an individual. BY and CS had the greatest CNVR diversity, with 835 and 820 unique CNVRs amounting to 13.8 Mb and 13.6 Mb respectively, as compared to 152 and 174 unique CNVRs comprising 0.6 Mb and 0.7 Mb in WL and WR. In addition, 160 CNVRs located on chrUn covered 1.5 Mb of genome sequence and may be copy number variable between individuals. Although we employed stringent quality control for those regions, candidate CNVRs on chrUn were worth a thorough study owing to the shorter length of the chrUn contigs and mapping ambiguity of chrUn sequence reads.

**Figure 1.**
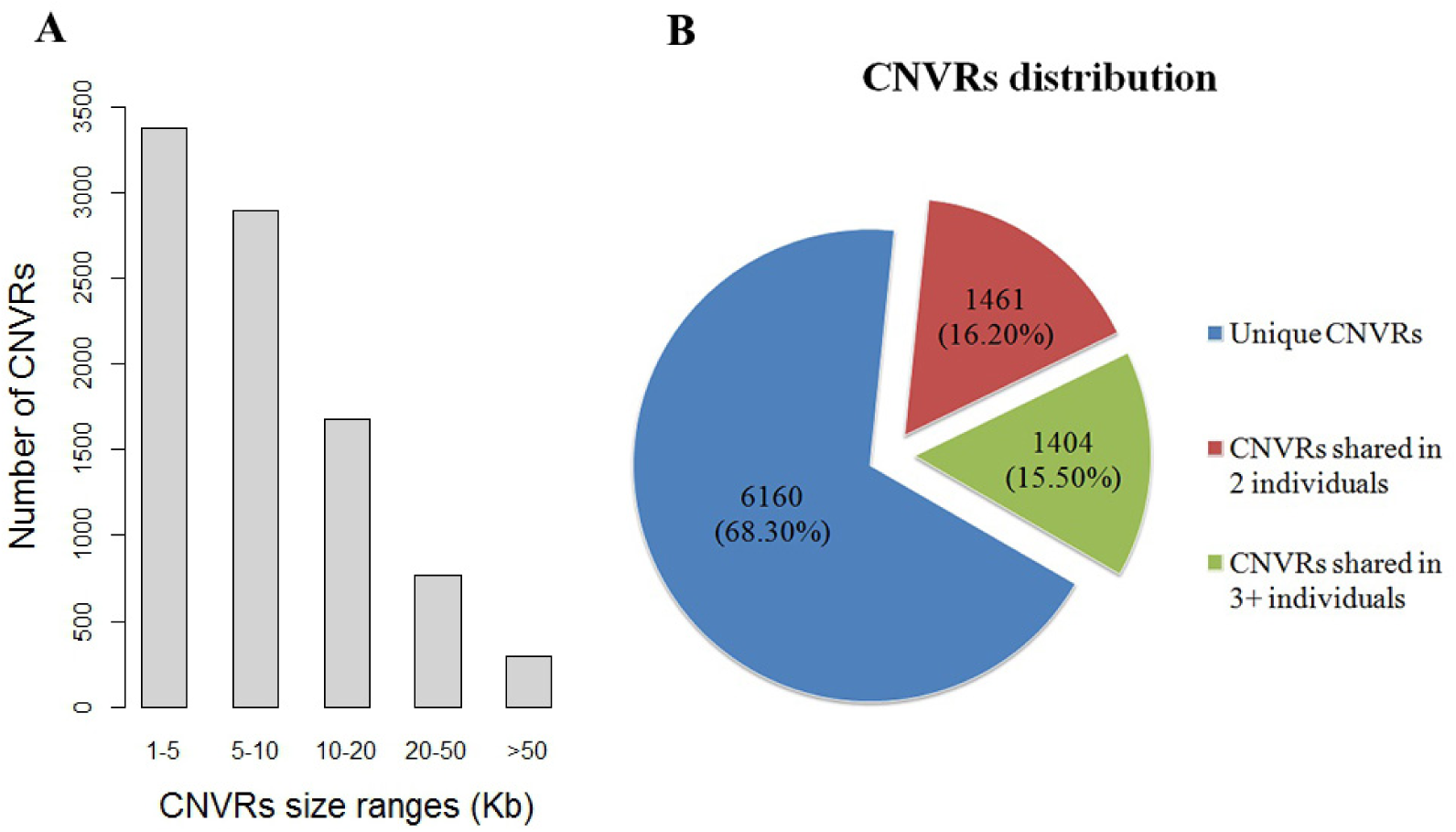
The length and frequency distribution of CNVRs. **(A)** Most CNVRs are shorter than 10 kb. **(B)** 6,CNVRs (68.30%) events occur in only one individual.

### Experimental validation

The copy number value of diploid regions in autosomes theoretically equals to two, so we could test the potential for CNVnator to generate false positive results by evaluating these two copies regions. For all 12 individuals, we selected all 5 kb non-overlapping windows in autosomes and excluded all windows intersecting with predicted CNVs and gaps, and then estimated their average CN. The average CN and STDEV per individual was 2.077 ± 0.291, varied from 2.041 ± 0.226 in WR to 2.104 ± 0.299 in RJF, showing low variability within the predicted neutral regions. Further, to validate sequencing-based CNV predictions, we carried out two independent experiments including aCGH and qPCR as two traditional CNV detection approaches to compare with computational predictions. We performed 11 pairwise aCGH experiments using RJF as the reference for all experiments and all others as test samples. Considering that we estimated CN of selected individuals with respect to reference genome which cannot be used for the aCGH reference sample, we calculated the predicted log_2_ CN ratios for the 11 aforementioned individuals against RJF based on computational copy number estimates to make the CN values comparable with the aCGH results, which was designated as digital aCGH approach (Sudmant et al. 2010). We first split the predicted overlapping CNVs from test samples and RJF into non-overlapping segments and estimated CN of each segment for each of the two samples, and divided the segment CN of test sample by RJF and calculated log_2_ CN ratios as digital aCGH values. Then we compared the digital values with aCGH probe log_2_ ratios which were defined as the average of all probes log_2_ ratio values in corresponding segments. We performed a simple linear regression analysis to explain the correlation between two values. Pearson correlation values (r) ranged from 0.395 in SK to 0.740 in LX among all 11 individuals (**Figure 2** and **Supplementary Figure S2**), and eight of which were greater than 0.600. BY (0.459), SK (0.395) and WR (0.477) showed lower correlation less than 0.500 compared with other individuals larger than 0.600, we found the mean of all probes log_2_ ratio values in the three aforementioned individuals were 1.05, 0.85 and 1.05 respectively, and were larger than the value of others which were close to zero.

**Figure 2.**
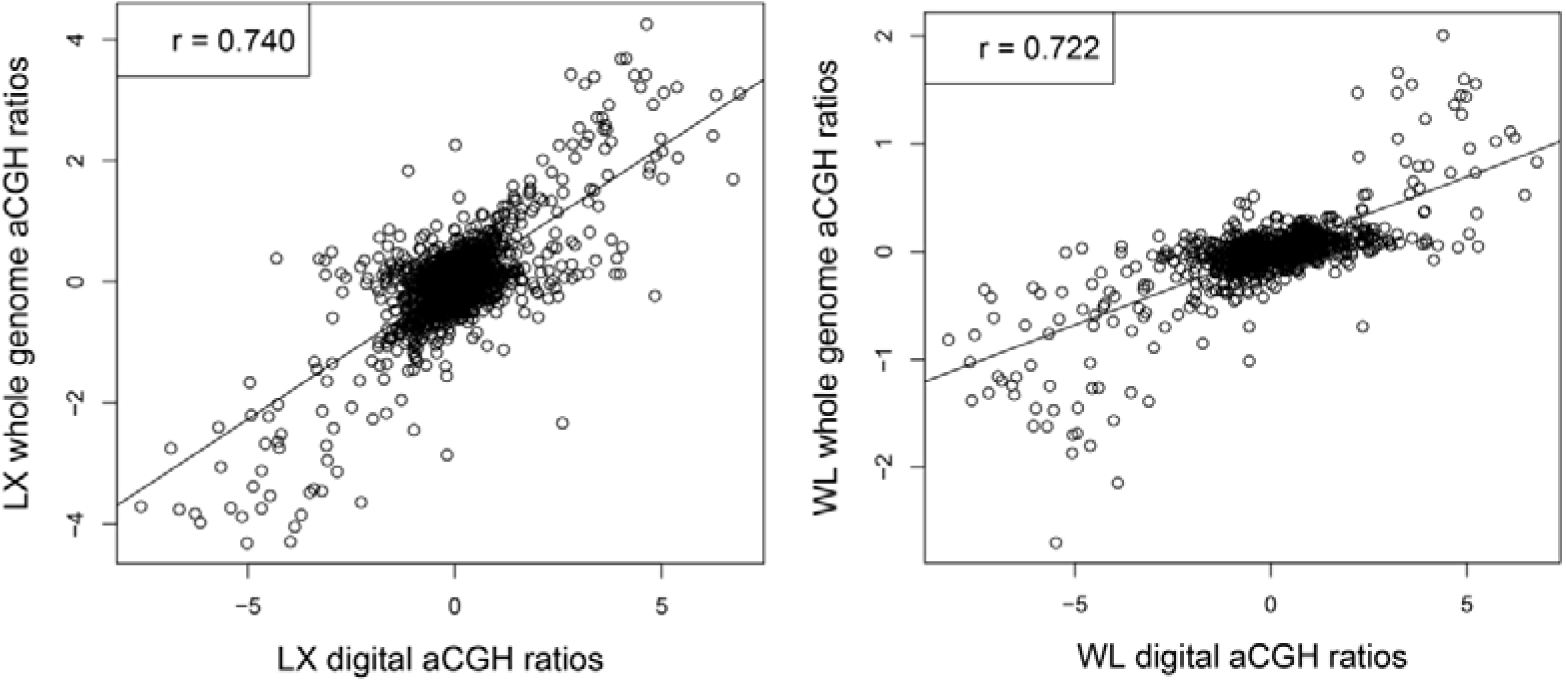
Correlation between digital aCGH and whole genome aCGH among Luxi Game and White Leghorn compared with Red Jungle Fowl (RJF). Digital aCGH are estimated using calculated logCN ratios in which CN are estimated for identified CNVs segments of two individuals and divided by the corresponding CN of RJF. RJF is selected as the reference sample in each aCGH experiment, and aCGH values are defined as the average of all probes logratio values in the same segments of digital aCGH.

In addition, we chose to investigate 15 predicted CNVRs representing different types and frequencies, and tested all 12 samples for each CNVR. Two distinct pairs of primers were designed for each predicted CNVR (**Supplementary Table S3**). The proportions of confirmed positive samples (positive predictive value) varied from 50% to 100%, with an average of 91.71%. However, some negative samples were also confirmed to contain CNV, and the false negative rate varied from 0 to 60%, with an average of 22.43%. We illustrated the qPCR results for three confirmed CNVRs of different types (gain, loss and both) (**Supplementary Figure S3**).

### Copy number polymorphic genes

We obtained 5,924 non-redundant RefSeq gene transcripts retrieved from the UCSC Genome browser and identified copy number polymorphic genes in different individuals through estimating the copy number of each gene by CNVnator. A total of 2,182 genes (36.8%) overlapped with 2,188 predicted CNVRs (24.2%), while the other 3,742 (63.2%) did not. Among them, 535 genes were found to be completely overlapped by CNVRs. The overlapping genes were found not to be highly duplicated sequences, and the maximum copy number is 12.0. We focused on the genes on the anchored chromosomes for the further analysis and discussion due to their clear chromosome locations. We identified the 25 most variable genes according to the STDEV of each gene CN in different individuals, and found that these genes were mainly involved in immune response and keratin formation (**Table 2** and **Supplementary Table S4**). The number of genes intersecting with putative CNVs in different individuals varied greatly, ranging from 154 in WR to 780 in BY, and only nine genes were shared by all surveyed individuals. Keratin gene families were detected to have large CN values and variances. Two significant CNVRs associated with dermal hyperpigmentation were located on chr20 at positions 11,217,001 to 11,272,200 (CNVR7984) and 11,651,801 to 11,822,900 (CNVR7990), which had already been described in detail in previous study (Dorshorst et al. 2011), and the distance between two loci was 379.6 kb. *SLMO2* and *TUBB1* as the candidate genes were completely covered by the first region which was predicted about twice as many copies of the region in DX and SK as in other individuals (**Figure 4a** **and Supplementary Figure S4a**). The major functional gene *EDN3* (endothelin 3) is not archived because predicted gene is not available for UCSC RefSeq database. We found that only BY had this CNVR while SK and DX as two typical breeds with dermal hyperpigmentation did not. So we further checked the raw results before removing CNVs overlapping with gaps. Two nearly identical CNVs were found, one at positions 11,111,501 to 11,238,600 in DX and the other at positions 11,111,401 to 11,238,900 in SK comprising two gaps larger than 100 bp, which were also confirmed by our whole genome aCGH (**Figure 4a** **and Supplementary Figure S4a**). The distance between the raw CNVR and the second region (CNVR7990) is 412.9 kb and almost perfectly supports the reported results (Dorshorst et al. 2011). Conversely, the first CNVR in BY (11,217,001 to 11,272,200) showing normal skin color does not contain *EDN3* gene (11,148,025 to 11,160,484), also provides evidence that copy number variable *EDN3* is the causal mutation resulting in dermal hyperpigmentation. Another previously identified CNVR involving *PRLR* (prolactin receptor) gene on chrZ (Elferink et al. 2008) was also detected in our study in which the CN of *PRLR* in WC and WL are twice as many as in other individuals. The sex-linked *K* allele containing two copies of *PRLR* in females is associated with late feathering and used widely for sexing hatchlings. Our sequencing-based and qPCR results showed that WC and WL should exhibit the late feathering phenotype, which is supported by actual phenotype record.

**Figure 3.**
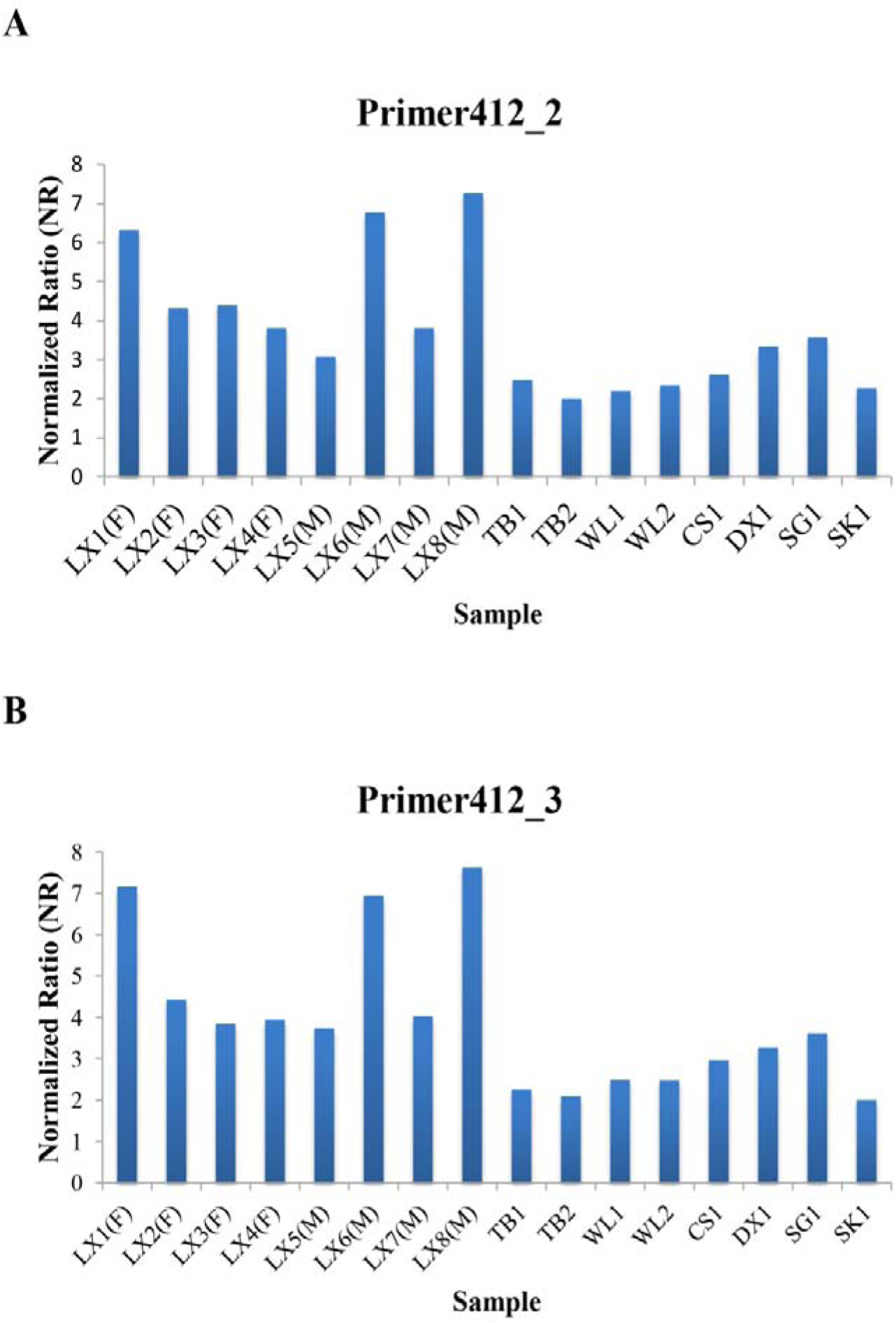
Validation of CNVR412 by qPCR in another 16 chickens. X-axis represents all 16 samples and Y-axis represents normalized ratios (NR) estimated by qPCR. NR around 2 indicates normal status (2 copies), NR around 0 or 1 indicates loss status (0 copies or 1 copy), and NR around 3 or more indicates gain status (3 or more copies).

**Figure 4.**
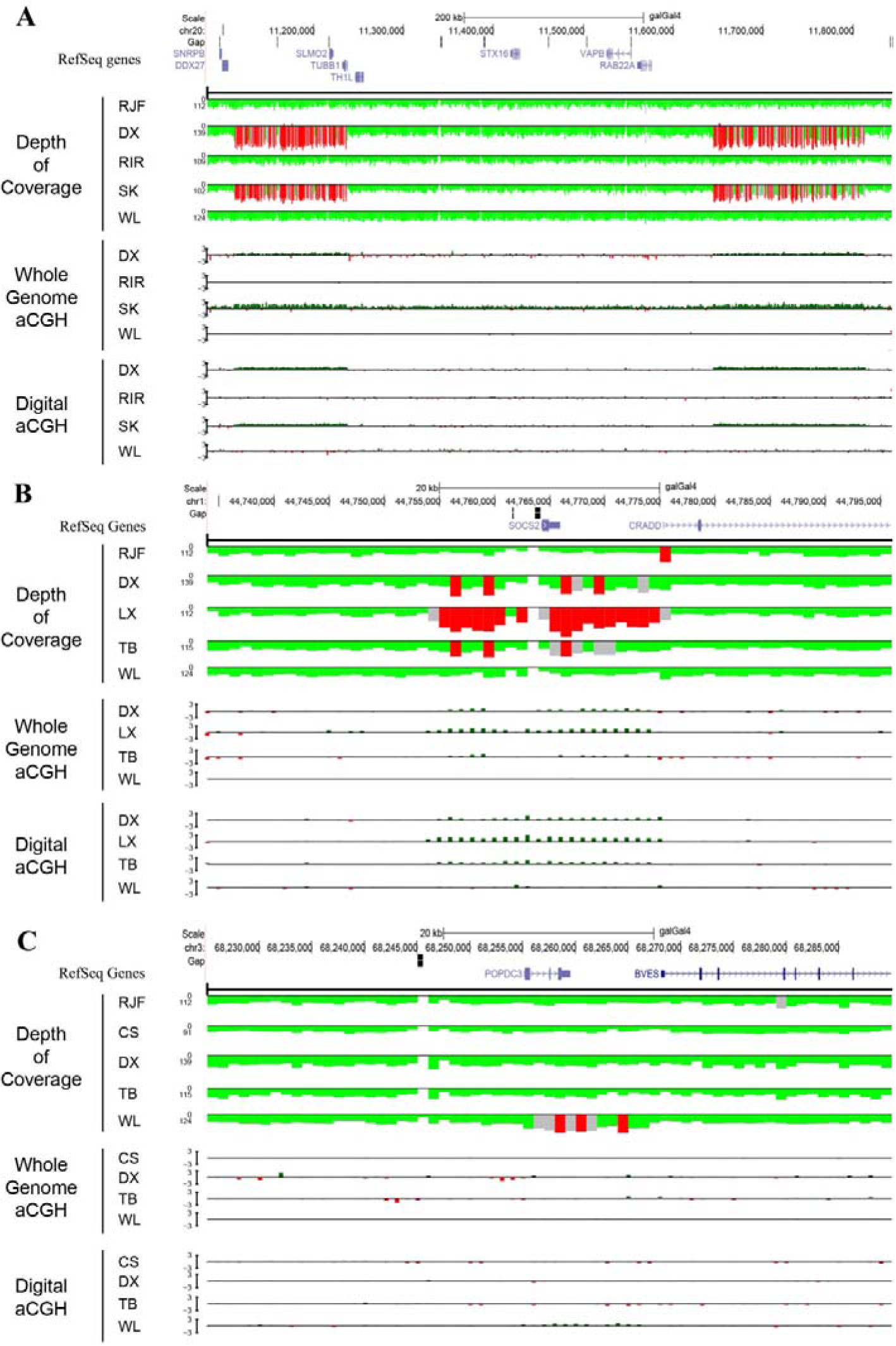
Read depth and digital aCGH predictions and whole-genome aCGH validations near preselected genetic loci for 5 representative chicken genomes. The uppermost gene image is generated with the UCSC Genome Browser (http://genome.ucsc.edu/) using galGal4 assembly. The track below the gene region is depth of coverage for all 5 individual genomes. Red indicates regions of excess read depth (> mean + 3 × STDEV), whereas gray indicates intermediate read depth (mean + 2 × STDEV < x < mean + 3 × STDEV), and green indicates normal read depth (mean±2 × STDEV). All read depth values based on 1 kb non-overlapping windows are corrected by GC content. Whole-genome aCGH and digital aCGH values are depicted as red-green histograms and correspond to a gain colored in green (> 0.5), a loss colored in red (< - 0.5) and normal status colored in gray (-0.5 < x < 0.5). **(A)** Two previous reported CNVs (chr20: 11,111,401-11,238,900 and chr20: 11,651,801-11,822,900) associated with dermal hyperpigmentation. The DX and SK genomes show two additional copies of these regions compared with RJF, and are validated by whole-genome aCGH. **(B)** A higher copy number increase for the *SOCS2* locus (chr1:44,764,280-44,765,955) is predicted in LX than in other individuals. (C) The *POPDC3* gene (chr3:68,255,196-68,259,535) is predicted duplication only in WL.

**Table 2.** Top copy number variable genes in chicken genomes

| Gene name | RefSeq accession | Gene size (bp) | Gene copy number estimates per individual<sup>a</sup> |  |  |  |  |  |  |  |  |  |  |  |
| --- | --- | --- | --- | --- | --- | --- | --- | --- | --- | --- | --- | --- | --- | --- |
|  |  |  | BY | CS | DX | LX | RIR | RJF | SG | SK | TB | WC | WL | WR |
| *LOC418424* | NM_001030786 | 7,212 | 2.8 | 3.5 | 2.4 | 4.3 | 4.0 | 3.2 | 6.3 | 4.6 | 8.5 | 3.1 | 2.6 | 2.0 |
| *LOC425362* | NM_001277974 | 648 | 4.0 | 2.9 | 3.4 | 4.2 | 2.5 | 6.0 | 7.1 | 4.1 | 5.2 | 3.0 | 1.7 | 2.9 |
| *C20H20ORF111* | NM_001029981 | 7,087 | 9.4 | 11.9 | 7.2 | 7.1 | 8.6 | 9.4 | 9.0 | 12.0 | 10.1 | 9.1 | 9.8 | 10.1 |
| *LOC100859427* | NM_001277807 | 1,118 | 4.7 | 3.5 | 3.0 | 3.6 | 2.8 | 4.1 | 3.1 | 7.9 | 4.0 | 5.1 | 2.9 | 2.8 |
| *SOCS2* | NM_204540 | 1,676 | 1.6 | 1.8 | 3.0 | 6.4 | 2.0 | 1.6 | 1.4 | 1.3 | 3.6 | 2.4 | 1.5 | 1.7 |
| *GBP* | NM_204652 | 2,531 | 1.4 | 0.2 | 4.1 | 2.3 | 2.1 | 2.5 | 3.1 | 4.4 | 4.5 | 3.1 | 2.0 | 3.6 |
| *PTRH2* | NM_001040413 | 1,161 | 1.6 | 3.2 | 1.1 | 1.3 | 3.6 | 1.8 | 1.4 | 1.2 | 5.1 | 3.2 | 1.5 | 1.9 |
| *LOC426914* | NM_001277964 | 981 | 4.9 | 2.7 | 3.2 | 3.9 | 2.2 | 4.0 | 1.9 | 5.6 | 4.2 | 4.4 | 2.4 | 3.0 |
| *PRSS2* | NM_205384 | 2,812 | 2.8 | 0.6 | 3.7 | 3.9 | 2.8 | 3.1 | 2.2 | 2.5 | 3.1 | 2.3 | 0.9 | 3.2 |
| *LOC100859722* | NM_001277975 | 660 | 2.6 | 3.1 | 3.6 | 4.2 | 5.1 | 5.2 | 4.1 | 2.2 | 3.6 | 4.0 | 4.8 | 4.4 |
| *LOC100859616* | NM_001277973 | 749 | 1.5 | 1.4 | 2.4 | 2.8 | 2.9 | 1.5 | 2.1 | 3.7 | 2.1 | 3.5 | 3.6 | 4.1 |
| *CD8A* | NM_001048080 | 3,183 | 3.5 | 3.6 | 5.2 | 2.5 | 2.7 | 2.1 | 3.9 | 2.5 | 2.5 | 3.0 | 2.4 | 3.0 |
| *LOC425137* | NM_001278080 | 5,930 | 3.0 | 3.7 | 2.9 | 1.9 | 3.8 | 2.6 | 1.7 | 3.4 | 3.2 | 4.9 | 2.9 | 2.5 |
| *LOC431317* | NM_001277978 | 586 | 1.6 | 2.8 | 2.1 | 2.6 | 4.0 | 3.9 | 3.5 | 1.3 | 2.2 | 3.1 | 3.0 | 2.9 |
| *SLMO2* | NM_001030866 | 4,953 | 2.3 | 2.0 | 4.0 | 2.2 | 2.1 | 2.4 | 2.1 | 4.5 | 2.5 | 2.2 | 2.0 | 1.9 |
| *TUBB1* | NM_205445 | 5,018 | 2.7 | 2.3 | 4.2 | 2.2 | 2.1 | 1.9 | 2.1 | 4.4 | 2.2 | 2.5 | 2.1 | 1.9 |
| *LIMS1* | NM_001001766 | 13,454 | 3.9 | 4.7 | 5.2 | 5.1 | 4.6 | 3.4 | 4.5 | 4.2 | 5.3 | 4.7 | 2.4 | 4.7 |
| *LOC770639* | NM_001277770 | 734 | 2.5 | 2.8 | 3.1 | 3.7 | 3.8 | 1.5 | 3.7 | 2.7 | 3.2 | 4.0 | 2.9 | 4.4 |
| *ZNF692* | NM_001099356 | 6,224 | 2.2 | 4.2 | 1.3 | 2.2 | 1.4 | 1.6 | 2.1 | 1.6 | 1.8 | 1.9 | 2.0 | 2.0 |
| *RFT1* | NM_001142872 | 10,993 | 2.1 | 2.2 | 1.9 | 2.1 | 3.6 | 2.0 | 1.9 | 2.1 | 4.1 | 3.0 | 2.0 | 1.9 |
| *LOC431316* | NM_001277972 | 741 | 1.6 | 2.3 | 1.4 | 1.9 | 3.7 | 2.8 | 2.4 | 1.4 | 2.2 | 3.2 | 3.0 | 2.1 |
| *LOC693258* | NM_001044681 | 720 | 2.8 | 4.2 | 2.5 | 3.4 | 4.0 | 3.5 | 2.4 | 2.3 | 2.6 | 3.1 | 2.3 | 2.3 |
| *LOC100859586* | NM_001277977 | 828 | 1.6 | 2.4 | 2.7 | 3.0 | 2.4 | 2.0 | 1.4 | 1.8 | 2.5 | 2.6 | 3.5 | 3.2 |
| *SOX3* | NM_204195 | 1,823 | 1.0 | 2.4 | 1.5 | 2.0 | 2.5 | 3.2 | 2.7 | 1.9 | 1.6 | 1.9 | 2.2 | 3.1 |
| *CD8A* | NM_205235 | 12,042 | 3.9 | 3.6 | 4.7 | 2.5 | 2.7 | 2.8 | 2.8 | 3.4 | 3.0 | 3.5 | 3.0 | 3.6 |

<sup>a</sup>BY, Beijing You; CS, Cornish; DX, Dongxiang; LX, Luxi Game; RIR, Red Island Rhode; RJF, Red Jungle Fowl; SG, Shouguang; SK, Silkie; TB, Tibetan; WC, Wenchang; WL, White Leghorn; WR, White Plymouth Rock.

In addition, we found some genes related to the host immune and inflammatory response. For example, *CD8A*, *FZD6*, *LIMS1*, *TNFSF13B* and some MHC genes associated with Marek’s disease (MD) were found to have CNVR overlaps, and the same case with genes for avian resistance to bacteria, such as *CDH13* and *CALM1*. *SOCS2* involving in the regulation of bone growth and density was predicted to have the largest CN values in LX (n = 6.4), while DX (n = 3.0) and TB (n = 3.6) also showed the duplicated sequence compared with the neutral of the other individuals in the loci (**Figure 4b** **and Supplementary Figure S4b**). LX represents a characteristic breed for cockfighting in which bone strength is an essential feature for selection. To validate the highly duplicated sequence (CNVR412) found only in LX, we selected another 16 individuals, *i.e.*, eight LX (four males and four females) and other eight females consisting of one CS, one DX, one SG, one SK, two TB and two WL, to perform qPCR experiments using the same two pairs of primers listed in **Supplementary Table S3**. Two qPCR results demonstrated copy number estimate of almost each LX was larger than others (**Figure 3**), and the average copy number (5.0 and 5.2 for two pairs of primers, respectively) of all LX were significant larger than those (2.6 and 2.6) in other individuals using the two-sample t-test (*P*-value = 0.003 and 0.001). Additionally, other identified CNV-gene overlaps were detected to be potentially responsible for economic traits, as these genes were involved in lipid metabolism, muscle development and growth, and secretion process containing hormone, protein and biotin. For example, our results suggested higher copy number for the *POPDC3* gene in WL (n = 4.2) than in the other 11 genomes (n = 2.3) (**Figure 4c** **and Supplementary Figure S4c**). Similarly, the WL genome showed the greatest number of *AVR2* copies (n = 2.0) on chrZ compared with others (n = 1.1). Two promising genes involving in lipid metabolism, *AP2M1* and *LBFABP*, were identified as the largest copy number (n = 3.0 and 3.2) in meat-type chicken (CS) compared with those of all others.

### Heatmap analysis

We performed a hierarchical clustering heatmap analysis and generated heatmaps based on Pearson’s correlation using the CN values for selected gene loci, in order to infer the potential evolutionary history of some genes among 12 individuals. Two genes *SLMO2* and *TUBB1* in DX and SK, were found to be highly duplicated regions and the two individuals were clustered into one group (**Figure 5a**). Another promising gene *SOCS2* was also confirmed for the difference in copy number between LX and others (Figure 5b). Meanwhile, WL showed specific expansion in *POPDC3* locus and was split into a separate clade (**Figure 5c**).

**Figure 5.**
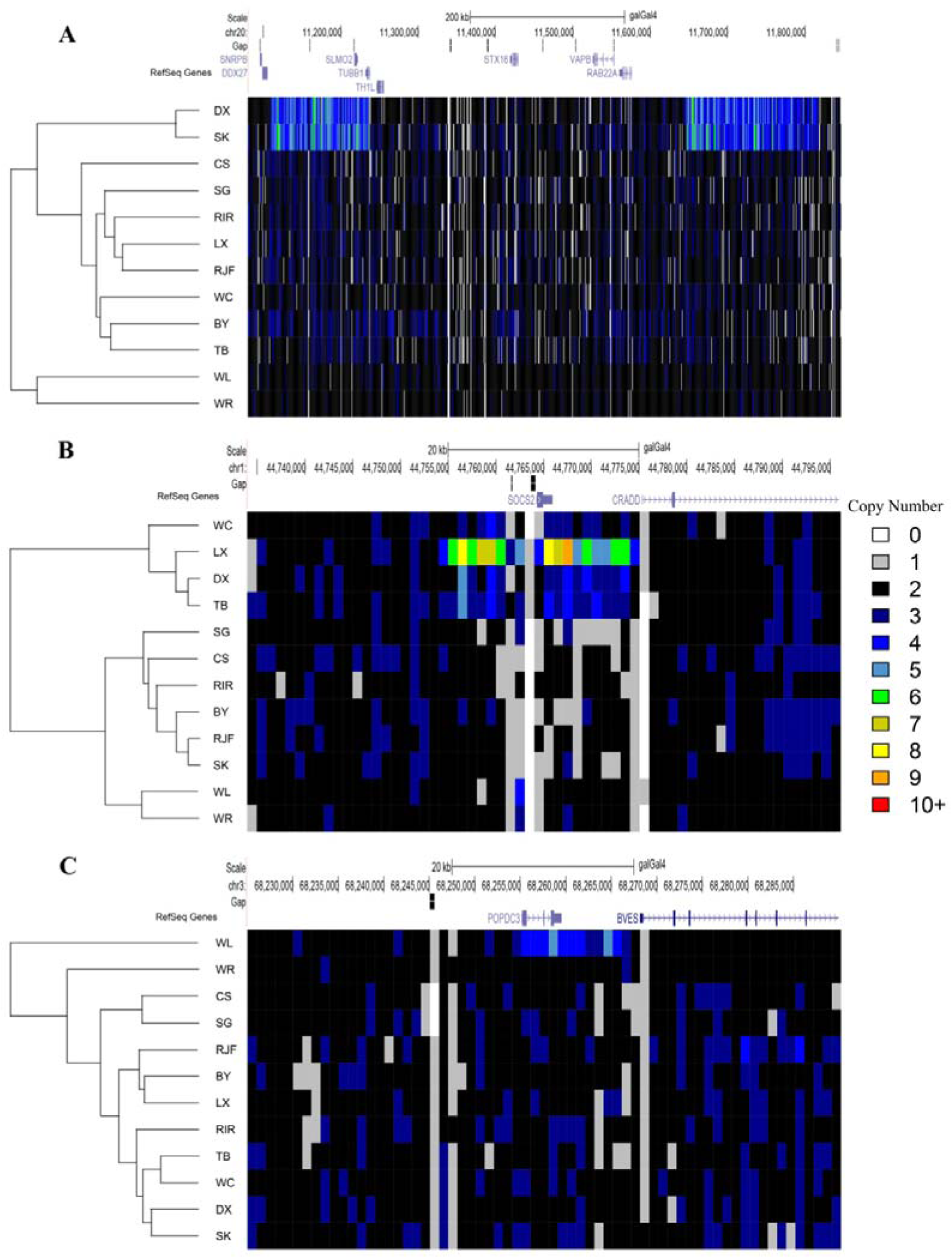
Hierarchical clustered heatmaps of preselected genetic loci for 12 chicken genomes. Every block in the heatmap indicates estimated CN values of kb non-overlapping windows in preselected region. These heatmaps are generated from hierarchical cluster analysis using Pearson’s correlation of the CN values. The colors for each bar denote different copy number (CN). **(A)** DX and SK which are predicted to be doubled within dermal hyperpigmentation loci are clustered together. **(B)** Upstream and downstream of *SOCS2* locus reveals higher CN values in DX, TB and WC especially LX. **(C)** WL shows specific expansion in *POPDC3* locus and is split into a separate clade.

### Gene content and QTL analysis of CNVRs

A total of 2,182 RefSeq genes overlapped putative CNVRs. Then, we performed gene ontology (GO) and Kyoto Encyclopedia of Genes and Genomes (KEGG) pathway analysis for these variable genes. The GO analysis revealed 641 GO terms, of which 157 were statistically significant after Benjamini correction (**Supplementary Table S5**). And GO terms showing significant enrichment were mainly involved in positive regulation of macromolecule metabolic process and gene expression, plasma membrane, protein localization, enzyme binding, response to oxidative stress and immune system development. The KEGG pathway analysis indicated that the variable genes were overrepresented in 11 pathways, but none of which was significant after Benjamini correction. According to our artificial QTL filtering criteria, we identified 595 high-confidence QTLs in total, of which 301 (50.6%) were found to overlap with 561 CNVRs (6.2%) (**Supplementary Table S6**). These QTLs were mainly involved in production and health traits, such as growth, body weight, abdominal fat weight, egg number and Marek’s disease-related traits.

## Discussion

This study performed genome-wide CNV detection, estimated absolute copy number values and constructed the first individualized chicken CNV map using NGS technology and RD method, which has advantages in both technology platform and genetic diversity compared with previous reports (Wang et al. 2012b; Crooijmans et al. 2013; Fan et al. 2013; Jia et al. 2013). CNV constitutes a major source of genetic variation that is complementary to SNP and could account for a substantial part of missing heritability (Manolio et al. 2009), because a significant fraction of CNVs fall in genomic regions not well covered by SNP arrays, especially SD regions lacking of sufficient probes (Campbell et al. 2011; Liu and Bickhart 2012). Most CNV studies to date have been discovery studies rather than association studies, mainly due to the limitations of CNV resolution and genotyping in each individual (McCarroll and Altshuler 2007). The high-resolution individualized chicken CNV map based on extensive genetic diversity not only enriches genetic variation database but also encourages the future development of assays for accurately genotyping CNVs, enabling systematic exploration about CNV association studies similar to SNPs. In future, integration of CNVs with SNPs may be an effective and promising way to elucidate the causes of complex diseases and traits (Stranger et al. 2007; Liu and Bickhart 2012).

The average number of putative CNVs per individual is 10 to 30 times more than that detected by previous aCGH studies (40 and 103 CNVs per individual; (Wang et al. 2012b; Crooijmans et al. 2013)) and four times more than our high-density aCGH results (391 CNVs per individual), and about 75% CNVs are smaller than 10 kb. It is mainly because most CNVs in genome are less than 10 kb in size, aCGH platforms with insufficient probes density have the limited capability of detecting them, whereas RD analysis is able to discover CNVs with a few hundred bases by increasing sequencing coverage (Abyzov et al. 2011). Additionally, the number of CNV events per individual in a recent report (4,419 CNVs; (Fan et al. 2013)) is larger than that in our results, owing to the difference between two CNV detection algorithms and post-filtering methods.

The number of CNVs and CNVRs even genes overlapping with CNVs in each individual varies greatly, and all individuals shares a small number of those, likely due to the distant relationship between 12 breeds for various breeding objectives. Out of all CNVRs, the percentage of CNVRs called in a single chicken breed is 68.3% and is similar to the other studies in chicken (71%, 73%, 64% and 62%; (Wang et al. 2010; Wang et al. 2012b; Luo et al. 2013; Tian et al. 2013)), while significantly higher than in human (49%; (Conrad et al. 2010)), cattle (32%; (Bickhart et al. 2012)) and dog (21%; (Berglund et al. 2012)). Because recombination rate is much higher in the chicken genome (2.5-21cM/Mb) compared with some mammalian such as human (1 cM/Mb) and mouse (0.5 cM/Mb) (Wong et al. 2004), and recombination-based mechanisms such as non-allelic homologous recombination (NAHR) are the major causes leading to CNVs (Munoz-Amatriain et al. 2013), we speculate that these specific CNVRs for a breed may be recent events and contribute to breed-specific phenotype and performance, and other CNVRs shared across breeds suggest their relative ancient origin or neutral evolutionary histories and seem to be fixed in all breeds. Meanwhile, the breed-specific CNVRs have smaller mean size because recent large scale variations may cause dysfunction and even be lethal (Conrad et al. 2006). Owing to only one individual per breed, a larger sample especially biological replicates within breed is crucial for validation study.

We find both maximum and mean copy number of duplicated sequences in chicken are less than those in mammalians (Alkan et al. 2009; Bickhart et al. 2012), which may be related to the relatively smaller genome size (only one third of a typical mammalian genome) and the lower repetitive DNA content in the chicken (Burt 2005). In addition, the covered sequence of gain CNVRs is larger than that of loss CNVRs because chromosomal deletion can lead to a variety of serious malformations and disorders and is subjected to purifying selection (Conrad et al. 2006; Freeman et al. 2006). In general, the length of chromosome is positively correlated with the number of CNVRs. The chr16 (a microchromosome) is found to have the second densest CNVRs, possibly owing to the highly variable major histocompatibility complex (MHC) regions and higher recombination rate, which result in the most genetic diversity of any chromosome (International Chicken Genome Sequencing Consortium 2004).

It is generally believed that the CN of neutral regions is between 1.5 and 2.5 (Abyzov et al. 2011) and the mean±2 × STDEV in our results corresponds closely to the theory, which demonstrates that CNVnator has efficient performance on CNV detection and CN estimation and can generate most reliable results. In addition, two independent validation experiments also suggest excellent accuracy and reliability of our predicted results. We first compare RD predicted CNVs with aCGH results, and the positive correlation between computational and experimental log_2_ CN ratios in our study is higher than the previous result (Bickhart et al. 2012), due to the two aCGH platforms with higher resolution for our analysis. The low correlation coefficients in BY, SK and WR may disclose certain experimental noises and biases resulting in misgenotyping in corresponding aCGH experiments (Liu and Bickhart 2012), particularly high-frequency duplications and rare deletions (Conrad et al. 2010; Abyzov et al. 2011). We then perform quantitative PCR for 15 randomly chosen CNVRs. The average of positive predicted value of the 15 validated CNVRs was 91.71%, similar to the results of previous reports in animals (Wang et al. 2012a; Jiang et al. 2013; Tian et al. 2013), suggesting that most of positive samples detected by sequencing-based are highly consistent with the qPCR experiments. Whereas we also estimate the false negative error rates as it is a common problem in CNV detection (Nicholas et al. 2009; Wang et al. 2012a), the average percentage of false negative results for each CNVR is 22.43%. This result may be due to the fact that we apply stringent criteria of CNV detection in order to minimize the false positive rate, while it also simultaneously results in possible increase in false negative rate.

Our results showed that 36.8% RefSeq genes intersected with 24.2% predicted CNVRs. It is probable that CNVs are located preferably in gene-poor regions (gene deserts and devoid of known regulatory elements), especially deletions (Conrad et al. 2006; Freeman et al. 2006), because gene-rich CNVs are more likely to be pathogenic than gene-poor CNVs and these deleterious CNVs are removed by purifying selection (Conrad et al. 2006; Lee et al. 2007). Meanwhile, the maximum CN of all genes overlapping with CNVs is 12.0, suggesting again that chicken genome has lower repetitive DNA content (Burt 2005). It is noted that some highly duplicated genes, especially nine out of the 25 most variable genes, belong to four keratin subfamilies (claw, feather, feather-like and scale) in chicken. In birds, skin appendages such as claws, scales, beaks and feathers are composed of beta (β) keratins and can prevent water loss and provide a barrier between the organism and its environment (Greenwold and Sawyer 2010), and the avian keratin genes are significant over-represented with respect to mammals (International Chicken Genome Sequencing Consortium 2004; Crooijmans et al. 2013). High CN keratin genes suggest the scenario for the evolution of the β-keratin gene family through gene duplication and divergence for their adaptive benefits (Zhang et al. 2009; Greenwold and Sawyer 2010). Additionally, the four subfamilies of β-keratin genes form a cluster on chr25 which is one of the more GC-rich chromosomes and contains a relatively larger number of minisatellites (Greenwold and Sawyer 2010), which also result in high copy number of genes.

CNV is a significant source of genetic variation accounting for disease and phenotypic diversity, due to the duplication or deletion of covered genes or regulation elements (Zhang et al. 2009), which are major forces of evolutionary innovation (Wapinski et al. 2007). Hierarchical clustering analysis of animals based on CN content within given locus could bring similar individuals during evolution into the same group and reveal the evolutionary relation shown by the heatmap. For example, a hierarchical clustering of CN values within *SLMO2* and *TUBB1* loci group DX and SK together, and both of which are distributed in the Jiangxi province of China, suggesting that DX and SK may have a close evolutionary relationship evolving from a common ancestor or purposely bred dermal hyperpigmentation into different strains. We detected CN differences for several interesting genes related to specific phenotypes among the surveyed individuals. For example, the *SOCS2* (suppressor of cytokine signaling 2) is a member of the suppressor of cytokine signaling family, the related proteins are implicated in the negative regulation of cytokine action through inhibition of the JAK/STAT pathway (Janus kinase/signal transducers and activators of transcription) (Metcalf et al. 2000). Dual x-ray absorptiometry (DXA) analysis demonstrated that *SOCS2* inactivation resulted in reduced trabecular and cortical volumetric bone mineral density (BMD) in *SOCS2*-deficient mice (Lorentzon et al. 2005). We find that *SOCS2* has the highest CN (n = 6.4) in LX than in the other individuals, which is particularly interesting as the LX is known for the cockfighting in which chickens with higher BMD have advantage over others. The gene expansions are also supported by heatmap. Additional qPCR experiments in other 16 individuals reveal that the increased copy number of *SOCS2* in LX is larger than others. We suspect that the copy number polymorphic locus is almost ubiquitous in chickens, but the particularly high gene duplication in LX may be as a result of the genetic effect of long-term artificial selection such as crossing between individuals with stronger bone.

Additionally, both the copy number estimates of *POPDC3* (popeye domain containing 3) and *AVR2* (avidin related protein 2) in WL were found to be about twice as many copies as other individuals. We draw a heatmap for *POPDC3* in WL to visualize specific gene duplication and clustering feature. The *POPDC3* gene belongs to Popeye family encoding proteins with three potential transmembrane domains with a high degree of sequence conservation, and is preferentially expressed in heart and skeletal muscle cells as well as smooth muscle cells (Brand 2005). It had been reported that the expression of two Popeye family members was upregulated in uterus of pregnant mice (Andree et al. 2000). Uterus has been thought to be an organ composed of smooth muscle and containing the shell gland in favor of depositing eggshell (Hincke et al. 2012), and duplication in *POPDC3* gene may facilitate myometrium maturation and labor as well as uterine fluid secretion during the egg laying period. Of the *AVR2* gene products, avidin is known to be the operational biotin-harvester produced in the oviducts of birds and deposited in the avian egg-white, comprising approximately 0.05% of the total protein in chicken egg-white. The function of *AVR2* has been postulated to be implicated in inflammation response in the manner of an antibiotic (Hytonen et al. 2005). WL is the most prolific egg laying chicken due to the fact that it has been extensively bred for egg production, thus the oviduct and uterus, serving as two important parts of the reproductive organs, are always in highly active state, and copy number increase at these loci related to laying may reveal important differences in abilities like protein secretion and eggshell formation between WL and other breeds.

Meat production is also a trait of economic importance. CS is a commonly used breed in the chicken meat industry and is found to have the largest CN of the *AP2M1* (adaptor-related protein complex 2, mu 1 subunit) and *LBFABP* (liver basic fatty acid binding protein) genes which are related to lipid metabolism and transport. *AP2M1* has been shown in microarray experiments to have higher expression in persons who fail to control their weight after weight reduction (Marquez-Quinones et al. 2010). Moreover, *LBFABP* is a member of the fatty acid-binding proteins (FABPs) family and expressed only in the liver playing a major role in lipid metabolism. It had been reported that feeding simulation was the primary factor increasing the expression of *LBFABP* gene (Murai et al. 2009). Duplication of the *AP2M1* and *LBFABP* locus in CS could potentially increase their expression, and may be associated with fatty acid utilization and weight gain.

Our findings suggest that many potential CNV-gene overlaps, like *CD8A*, *BF2* and *CALM1*, are associated with diseases susceptibility and resistance (Liaw et al. 2007; Goto et al. 2009; Connell et al. 2013), and also prove the two previous copy number variable genes involving in MD disease, namely *FZD6* (frizzled family receptor 6) and *LIMS1* (LIM and senescent cell antigen-like domains 1) (Wang-Rodriguez et al. 2002; Chen et al. 2008; Luo et al. 2013). Genes intersecting with CNVRs may be important sources of disease and phenotypic diversity through reshaping gene structure and modulating gene expression (Zhang et al. 2009). Moreover, these enriched GO terms are involved in cellular regulation and structure as well as various binding functions, in which most genes may be haploinsufficient, and duplication of them could improve fitness through selection on increased dosage effects (Nguyen et al. 2006). It is notable that several GO terms related to stress and immune response are overrepresented, suggesting that the CN variable genes may influence the responses to environmental stimuli and provide the mutational flexibility to adapt rapidly to changing selective pressures due to the signatures of adaptive evolution (Gokcumen et al. 2011).

## Conclusions

In this study, we performed genome-wide CNV detection and absolute copy number estimates of corresponding genetic locus based on the whole genome sequencing data of 12 chickens abundant in genetic diversity, and constructed the highest-resolution individualized chicken CNV map so far. We identified a total of 9,025 CNVRs in all individuals. Validation of CNVRs by aCGH and qPCR produced a high rate of confirmation, suggesting sequencing-based method was more sensitive and efficient for CNV discovery and genotyping. We have detected 2,182 RefSeq genes as copy number variable among 12 individuals, including genes involved in well-known phenotypes such as dermal hyperpigmentation and late feathering. In addition, some novel genes like *POPDC3* and *LBFABP* covered by CNVs may play an important role in production traits, and highly duplicated *SOCS2* may serve as an excellent candidate for bone mineral density. Our study based on extensive genetic diversity lays the foundation for comprehensive understanding of copy number variation in chicken genome and is beneficial to future association studies between CNV and important traits of chickens.

## Methods

### Sample collection and sequencing

We selected a total of 12 female chickens from different types and genetic sources representing modern chicken populations with abundant genetic diversity, *i.e.*, a Red Jungle Fowl (RJF, the ancestor of domestic chickens), seven Chinese indigenous chickens including Beijing You (BY), Dongxiang (DX), Luxi Game (LX), Shouguang (SG), Silkie (SK), Tibetan (TB) and Wenchang (WC), and four commercial breeds including Cornish (CS), Rhode Island Red (RIR), White Leghorn (WL) and White Plymouth Rock (WR). The whole blood samples were collected from brachial veins of chickens by standard venepuncture along with regular quarantine inspection of the experimental station of China Agricultural University, and genomic DNA was isolated using standard phenol/chloroform extraction method. Whole genome sequencing for all 12 individuals was performed on the HiSeq 2000 system (Illumina Inc., San Diego, CA, USA). Two genomic DNA libraries of 500 bp insert size per individual were constructed and sequenced with 100 bp paired-end reads, and each library dataset was generated with a five-fold coverage depth. Library preparation and all Illumina runs were performed as the standard manufacturer’s protocols.

### Quality control and Sequence alignment

For ensuring high-quality data, we used NGS QC Toolkit with default parameters to perform quality control of raw sequencing data, mainly by removing low-quality reads and reads containing primer/adaptor contamination (Patel and Jain 2012). All high-quality Illumina sequence reads were aligned against the galGal4 as a reference source by using Burrows-Wheeler Aligner (BWA) program (Li and Durbin 2009) with default parameters. The assembly of the reference genome was retrieved from the UCSC website (http://hgdownload.soe.ucsc.edu/goldenPath/galGal4/bigZips/). The BWA aligned output format was set to SAM. During the construction of a genomic library, Illumina platform was likely to generate some duplicate reads named ‘PCR and optical duplicates’ which imposed significant impact on the downstream analysis. So we first used SAMtools (Li et al. 2009) to convert the .sam files of different libraries belonging to the same individual to .bam files and sort and merge them, followed by removal of potential PCR duplicates using Picard (http://picard.sourceforge.net/).

### CNV detection

Following the above filtering step, the resulting .bam files were utilized for calling and genotyping of CNVs, post-processing were performed using CNVnator software based on RD method as previously described (Abyzov et al. 2011). CNVnator firstly calculated the count of mapped reads within user specified non-overlapping bins of equal size as the RD signal, and then adjusted the signal in consideration of a correlation of RD signal and GC content of the underlying genomic sequence. The mean-shift algorithm was employed to segment the signal with presumably different underlying CNs. Putative CNVs were predicted by applying statistical significance tests to the segments. A more detailed description of method could be found at CNVnator paper (Abyzov et al. 2011). We ran CNVnator with a bin size of 100 bp for our data. CNV calls were filtered using stringent criteria including a P-value < 0.01 and a size > 1 kb, and calls with > 50% of q0 (zero mapping quality) reads within the CNV regions were removed (q0 filter), and calls overlapping with gaps which is larger than or equal to 5 bp in the reference genome were excluded from consideration. In unknown chromosomes (chrN_random and chrun_random in UCSC, chrUn), we controlled CNV size to be shorter than arbitrary 1/10 total length of respective contig for reliable CNV detection considering the percentage of CNV versus macrochomosomes in length is approximate to 10% and CNV should be much shorter than a contig. Meanwhile, we performed genotyping of all 5 kb non-overlapping windows which did not overlap with putative CNVs and gaps in autosomes.

### aCGH validation

Initially, NimbleGen whole genome tiling array used in our experiment was a custom-designed 3*1.4 M array based on galGal4 2011 build, which contained a total of 1,425,178 50-75mer probes with a mean and median interval of 734 bp and 700 bp. The DNA labeling (Cy3 for samples and Cy5 for references), array hybridization, data normalization and scanning analysis were performed by NimbleGen Systems Inc. (Madison, WI, USA). Image and segmentation analysis were performed using NimbleScan 2.5 (segMNT algorithm) with parameter preset by the manufacturer. However, there was some trouble during the NimbleGen aCGH experiments. Because none of results were obtained in three consecutive trials for CS, RIR and WL and this type of NimbleGen CGH array stopped production subsequently. Considering we only analyzed raw aCGH log_2_ ratio values instead of processed/normalized data, so we chose a similar Agilent custom-designed 1*1.0 M array (Agilent Technology Inc., CA, USA) with the mean and median probe spacing of 1,056 bp and 1,050 bp, respectively. And all data processing was performed in terms of standard Agilent procedure. In each aCGH experiment, we chose the RJF as the same reference sample.

### Quantitative PCR confirmation

We also performed qPCR confirmation of 15 CNVRs chosen from the CNVRs detected by CNVnator. Most chosen CNVRs have not been reported in previous studies and are also adjacent to annotated genes. Two distinct pairs of PCR primers were designed to target each CNV region using Primer5.0 software for the uncertainty in CNVR breakpoints. Furthermore, the UCSC In-Silico PCR tool was used for in silico analysis of primers specificity and sensitivity (Karolchik et al. 2008). PCCA which was previously identified as a non-CNV locus was chosen as a control region (Wang et al. 2010). Quality control of all primer sets were evaluated using an 8-point standard curve in duplicate to ensure the similar amplification efficiencies between target and control primers. All qPCR experiments were conducted on an ABI Prism 7500 sequence detection system (Applied Biosystems group) using SYBR green chemistry in triplicate reactions, each with a reaction volume of 15μl in a 96-well plate. The condition for thermal cycle was as follows: 1 cycle of pre-incubation at 50°C for 2 min and 95°C for 10 min, 40 cycles of amplification (95°C for 10 s and 60°C for 1 min). We used the formula 2^(1 - ΔΔCt)^ method to calculate the relative copy number for each test region by assuming that there were two copies of DNA in the control region. The cycle threshold (Ct) value of each test sample was normalized to the control region first, and then the ΔCt value was calculated between the test sample and a preselected reference sample predicted without CNV by CNVnator. The golden standard of each diploid CNVR was generally considered to have two copies for autosomes or one copy when the locus was on Z chromosome of a female in chickens.

### Gene contents and functional annotation

The RefSeq gene list was retrieved from the UCSC RefSeq database (Karolchik et al. 2008). All miRNA genes were excluded because the nucleotide sequences were too short to estimate reliable copy number. We analyzed the proportion of the RefSeq genes overlapping with putative CNVRs and performed CN estimates on all 5,927 non-redundant RefSeq gene transcripts. In addition, to provide insight into the functional enrichment of the RefSeq genes overlapping with CNVRs, we performed Gene Ontology (GO) functional annotation and Kyoto Encyclopedia of Genes and Genomes (KEGG) pathway analysis employing the web-accessible program DAVID (Huang da et al. 2009) and selecting the DAVID default population background which was appropriate for high-throughput studies in enrichment calculation. Statistical significance was accessed by using *P* value (P < 0.05) of a modified Fisher’s exact test and Benjamini correction for multiple testing. We also investigated the CNVRs identified in this study with the reported QTLs obtained from Chicken QTL database (Hu et al. 2013). We focused on the QTLs with confidence interval less than 10 Mb and considered those QTLs with overlapped confidence intervals greater than 50% as the same QTL (Jiang et al. 2013), because the QTL confidence intervals were too large to be used efficiently in post-processing.

### Heatmap hierarchical cluster analysis

We used the heatmap.2() function of the gplots package (http://cran.r-project.org/web/ packages/gplots/index.html) to generate heatmap figures. We first selected the specified regions extending 30 kb on each side of interesting genes and used the estimated CN values of 1 kb non-overlapping windows for each animal for post analysis, mainly considering that some regulatory elements may be included in the upstream or downstream of a gene. No reordering of those windows representing corresponding chromosome locations in heatmap was made for the sake of clarity. The Pearson’s correlation coefficient (1-r) of the CN values was used as a distance measure of the agglomerative hierarchical clustering with average linkage, and to generate hierarchical cluster dendrograms for each animal.

## Data access

All aCGH data have been submitted to the NCBI Gene Expression Omnibus (GEO) (http://www.ncbi.nlm.nih.gov/geo/) under accession number GSE54119.

## Acknowledgements

We thank Xiquan Zhang for sharing some samples. This work was funded in part by Programs for Changjiang Scholars and Innovative Research in University (IRT1191), and China Agriculture Research Systems (CARS-41).

## Authors’ contributions

N.Y. and L.Q. conceived and designed all experiments. G.Y., L.Q. and Y. Y. performed bioinformatics and statistical analysis with help from J.L., and carried out aCGH and qPCR experiments. G.X provided samples. G.Y. and L.Q. drafted the manuscript. N.Y. revised the paper. All authors read and approved the final manuscript.

## Supplementary figure legends

**Supplementary Figure S1.**
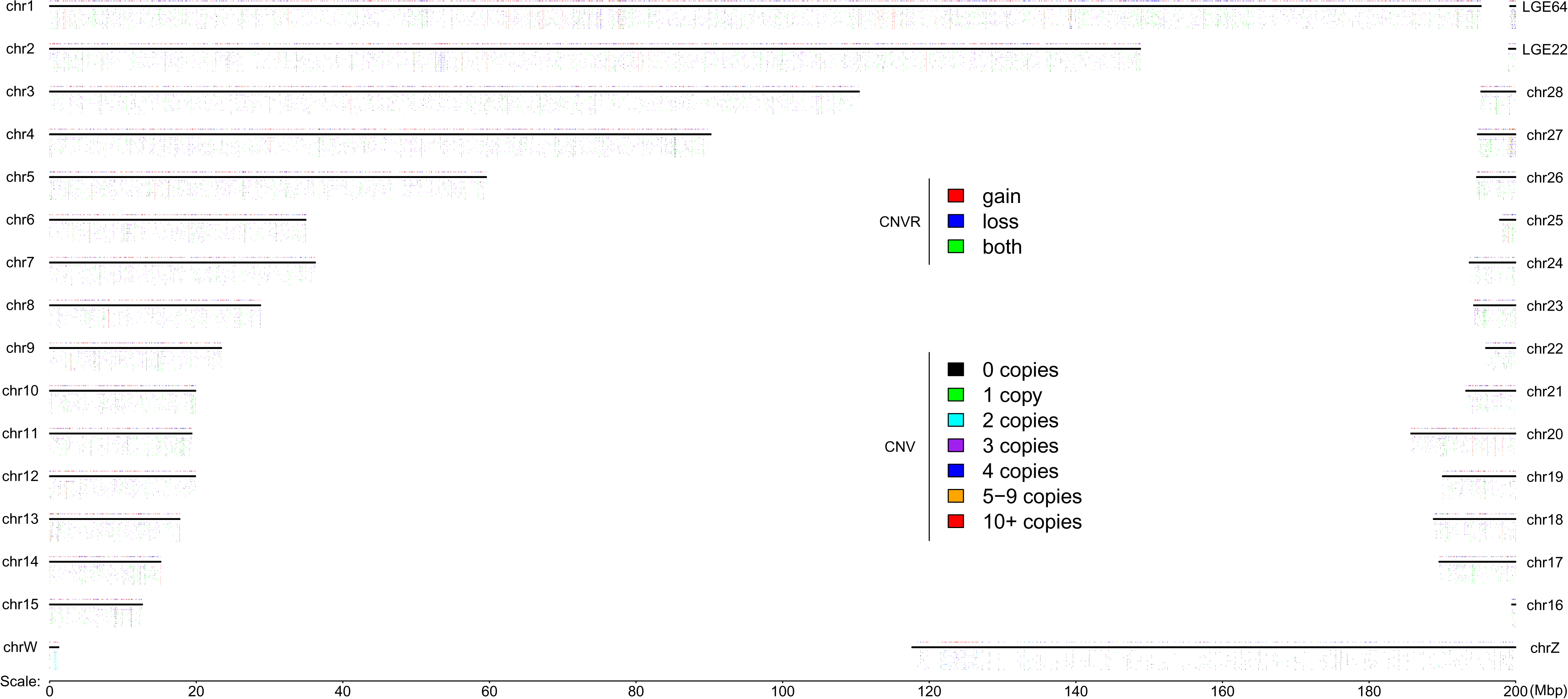
**Individualized chicken CNV map in the chicken genome.** The horizontal black lines represent the draft chicken genome (UCSC version galGal4). Tracks under the chromosomes indicate corresponding CNV status of all individuals kept in the alphabetical order from top to bottom, for BY, CS, DX, LX, RIR, RJF, SG, SK, TB, WC, WL and WR. Merged CNVRs from all individuals are depicted above chromosomes. The colors for each bar denote different copy number (CN) in CNV legend and different types of CNVRs. The downmost axis shows the chromosomes and CNVs coordinate. Left-hand chromosomes are ordered from left to right, and the right-hands are just reversed.

**Supplementary Figure S2.**
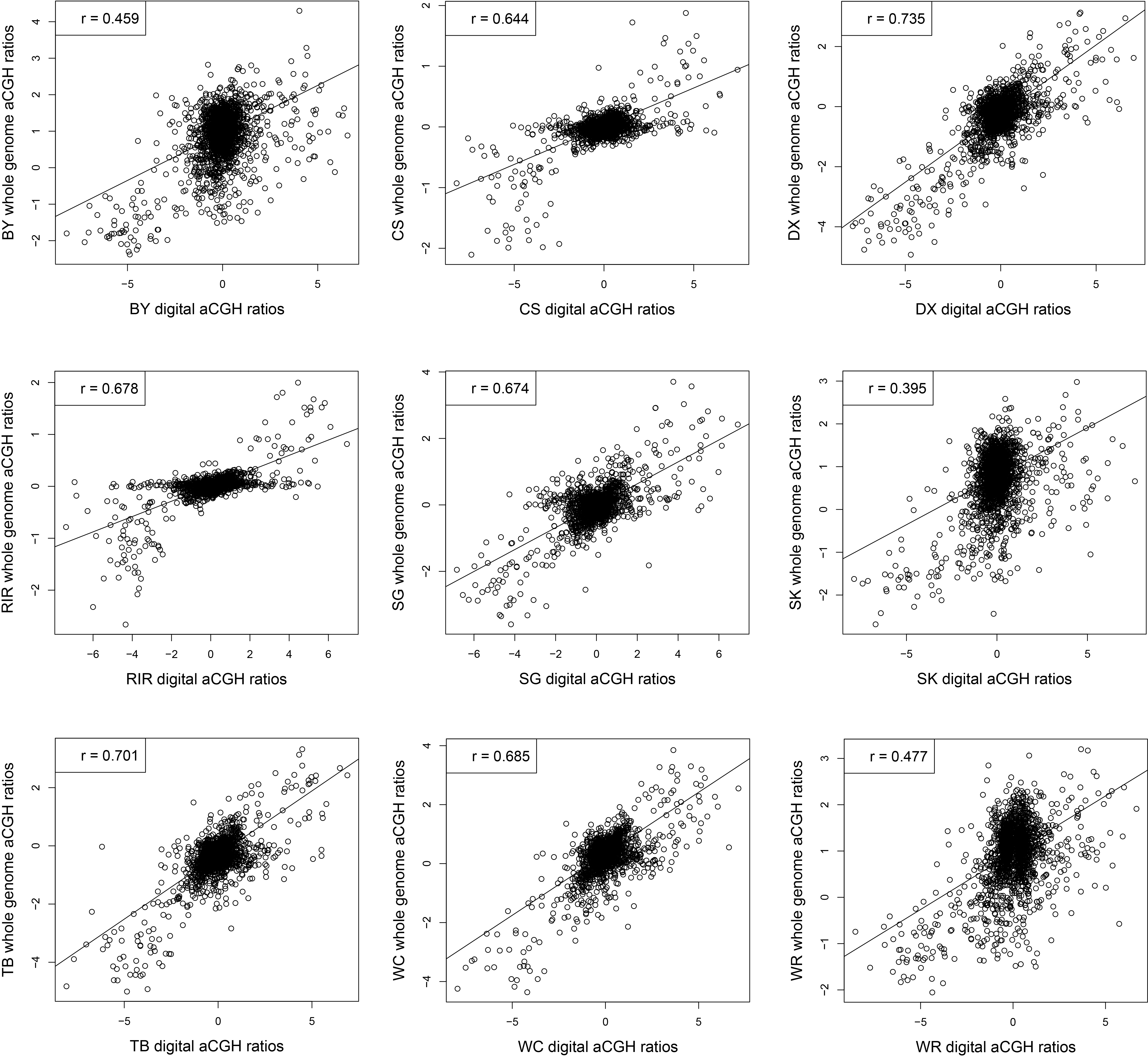
**Correlation between digital aCGH and whole-genome aCGH among nine individuals compared with Red Jungle Fowl (RJF).** Digital aCGH are estimated using calculated log_2_ CN ratios in which CN are estimated for identified CNVs segments of nine individuals and divided by the corresponding CN of RJF. RJF is selected as the reference sample in each aCGH experiment, and aCGH values are defined as the average of all probes log_2_ ratio values in the same segments of digital aCGH.

**Supplementary Figure S3.**
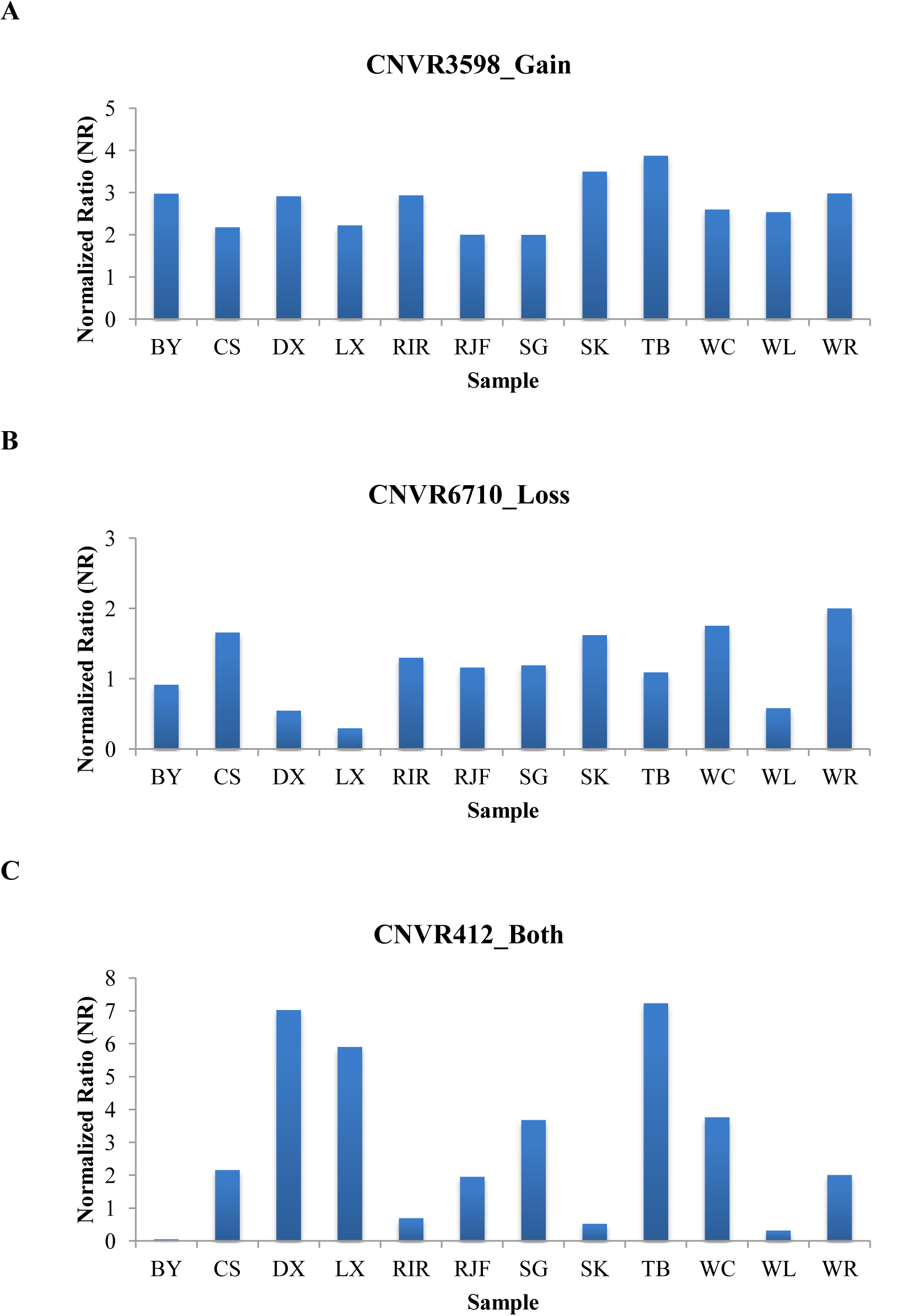
**Illustrating of qPCR confirmation results for three selected CNVRs of different types.** X-axis represents all 12 samples and Y-axis represents normalized ratios (NR) estimated by qPCR. NR around 2 indicates normal status (2 copies), NR around 0 or 1 indicates loss status (0 copies or 1 copy), and NR around 3 or more indicates gain status (3 or more copies). **(A)** Results for a gain status of CNVR3598. **(B)** Results for a loss status of CNVR6710. **(C)** Results for a both status of CNVR412.

**Supplementary Figure S4.**
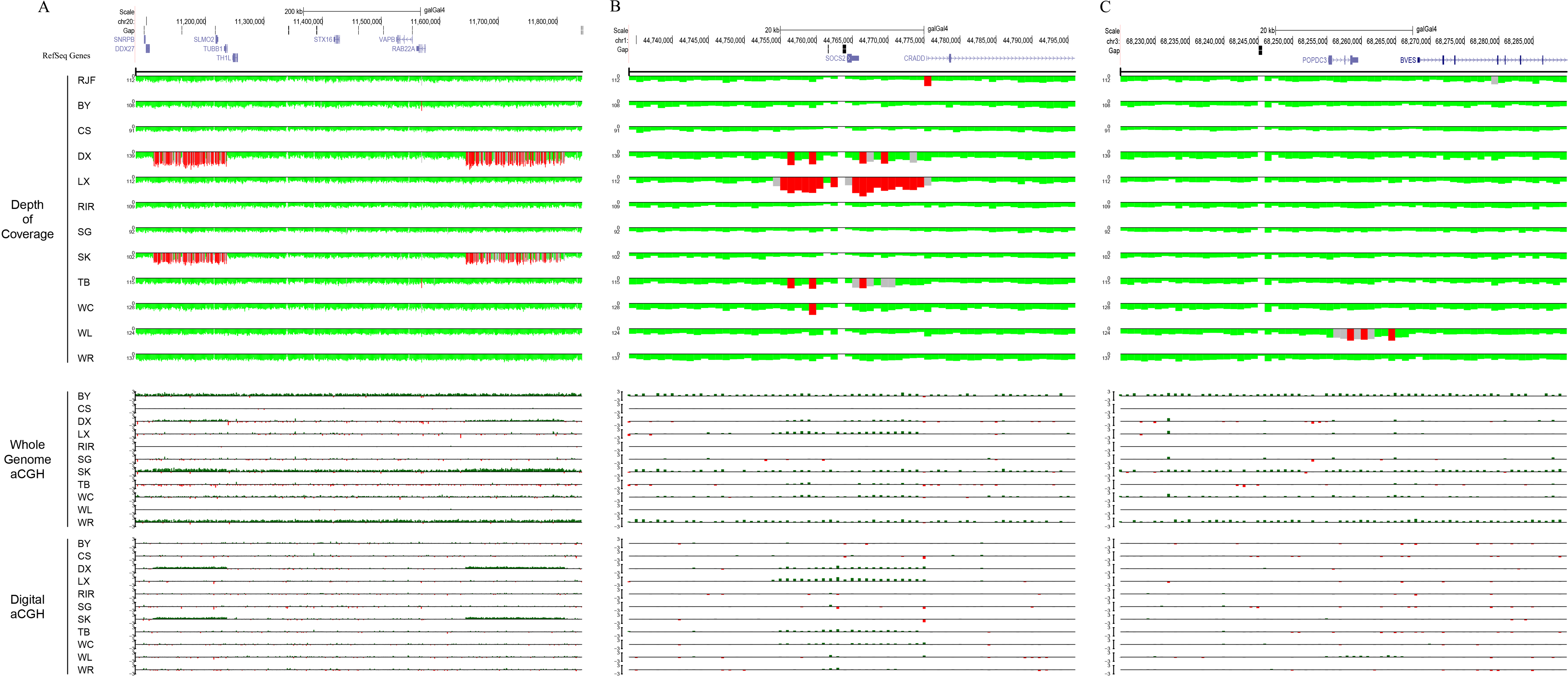
**Read depth and digital aCGH predictions and whole-genome aCGH validations near preselected genetic loci for 12 chicken genomes.** The uppermost gene image is generated with the UCSC Genome Browser (http://genome.ucsc.edu/) using galGal4 assembly. The track below the gene region is depth of coverage for all 12 individual genomes. Red indicates regions of excess read depth (> mean + 3 × STDEV), whereas gray indicates intermediate read depth (mean + 2 × STDEV < x < mean + 3 × STDEV), and green indicates normal read depth (mean ± 2×STDEV). All read depth values based on 1 kb non-overlapping windows are corrected by GC content. Whole-genome aCGH and digital aCGH values are depicted as red-green histograms and correspond to a gain colored in green (> 0.5), a loss colored in red (<-0.5) and normal status colored in gray (-0.5 < x < 0.5). **(A)** Two previous reported CNVs (chr20: 11,111,401-11,238,900 and chr20: 11,651,801-11,822,900) associated with dermal hyperpigmentation. The DX and SK genomes show two additional copies of these regions compared with RJF, and are validated by whole-genome aCGH. **(B)** A higher copy number increase for the *SOCS2* locus (chr1:44,764,280-44,765,955) is predicted in LX than in other individuals. **(C)** The POPDC3 gene (chr3:68,255,196-68,259,535) is predicted duplication only in WL.

## Supplementary tables

**Supplementary Table S1. Summary of identified CNVs and CNVRs in 12 chicken genomes.**

**Supplementary Table S2. General statistics of the CNVRs on each chromosome.**

**Supplementary Table S3. Primers information and confirmation results of the 15 chosen CNVRs by qPCR analysis.**

**Supplementary Table S4. The detailed features of RefSeq genes completely or partial overlapping with CNVRs.**

**Supplementary Table S5. Functional enrichment of GO and KEGG pathway analysis of RefSeq genes covered by CNVRs.**

**Supplementary Table S6. The overlap information of QTLs and CNVRs across the chicken genome.**

## References

Abyzov A, Urban AE, Snyder M, Gerstein M. 2011. CNVnator: an approach to discover, genotype, and characterize typical and atypical CNVs from family and population genome sequencing. Genome Res 21(6): 974–984.

Alkan C, Kidd JM, Marques-Bonet T, Aksay G, Antonacci F, Hormozdiari F, Kitzman JO, Baker C, Malig M, Mutlu O et al. 2009. Personalized copy number and segmental duplication maps using next-generation sequencing. Nat Genet 41(10): 1061–1067.

Altshuler DM, Gibbs RA, Peltonen L, Dermitzakis E, Schaffner SF, Yu F, Bonnen PE, de Bakker PI, Deloukas P, Gabriel SB et al. 2010. Integrating common and rare genetic variation in diverse human populations. Nature 467(7311): 52–58.

Andree B, Hillemann T, Kessler-Icekson G, Schmitt-John T, Jockusch H, Arnold HH, Brand T. 2000. Isolation and characterization of the novel popeye gene family expressed in skeletal muscle and heart. Dev Biol 223(2): 371–382.

Bentley DR Balasubramanian S Swerdlow HP Smith GP Milton J Brown CG Hall KP Evers DJ Barnes CL Bignell HR et al. 2008. Accurate whole human genome sequencing using reversible terminator chemistry. Nature 456(7218): 53–59.

Berglund J, Nevalainen EM, Molin AM, Perloski M, Andre C, Zody MC, Sharpe T, Hitte C, Lindblad-Toh K, Lohi H et al. 2012. Novel origins of copy number variation in the dog genome. Genome Biol 13(8): R73.

Bickhart DM, Hou Y, Schroeder SG, Alkan C, Cardone MF, Matukumalli LK, Song J, Schnabel RD, Ventura M, Taylor JF et al. 2012. Copy number variation of individual cattle genomes using next-generation sequencing. Genome Res 22(4): 778–790.

Brand T. 2005. The Popeye domain-containing gene family. Cell Biochem Biophys 43(1): 95–103.

Burt DW. 2005. Chicken genome: current status and future opportunities. Genome Res 15(12): 1692–1698.

Campbell CD, Sampas N, Tsalenko A, Sudmant PH, Kidd JM, Malig M, Vu TH, Vives L, Tsang P, Bruhn L et al. 2011. Population-genetic properties of differentiated human copy-number polymorphisms. Am J Hum Genet 88(3): 317–332.

Chen K, Tu Y, Zhang Y, Blair HC, Zhang L, Wu C. 2008. PINCH-regulates the ERK-Bim pathway and contributes to apoptosis resistance in cancer cells. J Biol Chem 283(5): 2508–2517.

Connell S, Meade KG, Allan B, Lloyd AT, Downing T, O’Farrelly C, Bradley DG. 2013. Genome-wide association analysis of avian resistance to Campylobacter jejuni colonization identifies risk locus spanning the CDH13 gene. G3(Bethesda) 3(5): 881–890.

Conrad DF, Andrews TD, Carter NP, Hurles ME, Pritchard JK. 2006. A high-resolution survey of deletion polymorphism in the human genome. Nat Genet 38(1): 75–81.

Conrad DF, Pinto D, Redon R, Feuk L, Gokcumen O, Zhang Y, Aerts J, Andrews TD, Barnes C, Campbell P et al. 2010. Origins and functional impact of copy number variation in the human genome. Nature 464(7289): 704–712.

Crooijmans RP, Fife MS, Fitzgerald TW, Strickland S, Cheng HH, Kaiser P, Redon R, Groenen MA. 2013. Large scale variation in DNA copy number in chicken breeds. BMC Genomics 14: 398.

Dorshorst B, Molin AM, Rubin CJ, Johansson AM, Stromstedt L, Pham MH, Chen CF, Hallbook F, Ashwell C, Andersson L. 2011. A complex genomic rearrangement involving the endothelin 3 locus causes dermal hyperpigmentation in the chicken. PLoS Genet 7(12): e1002412.

Elferink MG, Vallee AA, Jungerius AP, Crooijmans RP, Groenen MA. 2008. Partial duplication of the PRLR and SPEF2 genes at the late feathering locus in chicken. BMC Genomics 9: 391.

Fan WL, Ng CS, Chen CF, Lu MY, Chen YH, Liu CJ, Wu SM, Chen CK, Chen JJ, Mao CT et al. 2013. Genome-wide patterns of genetic variation in two domestic chickens. Genome Biol Evol 5(7): 1376–1392.

Freeman JL, Perry GH, Feuk L, Redon R, McCarroll SA, Altshuler DM, Aburatani H, Jones KW, Tyler-Smith C, Hurles ME et al. 2006. Copy number variation: new insights in genome diversity. Genome Res 16(8): 949–961.

Gokcumen O, Babb PL, Iskow RC, Zhu Q, Shi X, Mills RE, Ionita-Laza I, Vallender EJ, Clark AG, Johnson WE et al. 2011. Refinement of primate copy number variation hotspots identifies candidate genomic regions evolving under positive selection. Genome Biol 12(5): R52.

Goto RM, Wang Y, Taylor RL, Jr., Wakenell PS, Hosomichi K, Shiina T, Blackmore CS, Briles WE, Miller MM. 2009. BG1 has a major role in MHC-linked resistance to malignant lymphoma in the chicken. Proc Natl Acad Sci U S A 106(39): 16740–16745.

Greenwold MJ, Sawyer RH. 2010. Genomic organization and molecular phylogenies of the beta (beta) keratin multigene family in the chicken (Gallus gallus) and zebra finch (Taeniopygia guttata): implications for feather evolution. BMC Evol Biol 10: 148.

Griffin DK, Robertson LB, Tempest HG, Vignal A, Fillon V, Crooijmans RP, Groenen MA, Deryusheva S, Gaginskaya E, Carre W et al. 2008. Whole genome comparative studies between chicken and turkey and their implications for avian genome evolution. BMC Genomics 9: 168.

Hastings PJ, Ira G, Lupski JR. 2009. A microhomology-mediated break-induced replication model for the origin of human copy number variation. PLoS Genet 5(1): e1000327.

Henrichsen CN, Chaignat E, Reymond A. 2009. Copy number variants, diseases and gene expression. Hum Mol Genet 18(R1): R1–8.

Hincke MT, Nys Y, Gautron J, Mann K, Rodriguez-Navarro AB, McKee MD. 2012. The eggshell: structure, composition and mineralization. Front Biosci (Landmark Ed) 17: 1266–1280.

Hu ZL, Park CA, Wu XL, Reecy JM. 2013. Animal QTLdb: an improved database tool for livestock animal QTL/association data dissemination in the post-genome era. Nucleic Acids Res 41(Database issue): D871–879.

Huang da W, Sherman BT, Lempicki RA. 2009. Systematic and integrative analysis of large gene lists using DAVID bioinformatics resources. Nat Protoc 4(1): 44–57.

Hytonen VP, Maatta JA, Kidron H, Halling KK, Horha J, Kulomaa T, Nyholm TK, Johnson MS, Salminen TA, Kulomaa MS et al. 2005. Avidin related protein 2 shows unique structural and functional features among the avidin protein family. BMC Biotechnol 5: 28.

International Chicken Genome Sequencing Consortium. 2004. Sequence and comparative analysis of the chicken genome provide unique perspectives on vertebrate evolution. Nature 432(7018): 695–716.

Jia X, Chen S, Zhou H, Li D, Liu W, Yang N. 2013. Copy number variations identified in the chicken using a 60 K SNP BeadChip. Anim Genet 44(3): 276–284.

Jiang L, Jiang J, Yang J, Liu X, Wang J, Wang H, Ding X, Liu J, Zhang Q. 2013. Genome-wide detection of copy number variations using high-density SNP genotyping platforms in Holsteins. BMC Genomics 14: 131.

Karolchik D, Kuhn RM, Baertsch R, Barber GP, Clawson H, Diekhans M, Giardine B, Harte RA, Hinrichs AS, Hsu F et al. 2008. The UCSC Genome Browser Database: 2008 update. Nucleic Acids Res 36(Database issue): D773–779.

LaFramboise T. 2009. Single nucleotide polymorphism arrays: a decade of biological, computational and technological advances. Nucleic Acids Res 37(13): 4181–4193.

Lee C, Iafrate AJ, Brothman AR. 2007. Copy number variations and clinical cytogenetic diagnosis of constitutional disorders. Nat Genet 39(Suppl): S48–54.

Li H, Durbin R. 2009. Fast and accurate short read alignment with Burrows-Wheeler transform. Bioinformatics 25(14): 1754–1760.

Li H, Handsaker B, Wysoker A, Fennell T, Ruan J, Homer N, Marth G, Abecasis G, Durbin R. 2009. The Sequence Alignment/Map format and SAMtools. Bioinformatics 25(16): 2078–2079.

Liaw HJ, Chen WR, Huang YC, Tsai CW, Chang KC, Kuo CL. 2007. Genomic organization of the chicken CD8 locus reveals a novel family of immunoreceptor genes. J Immunol 178(5): 3023–3030.

Liu GE, Bickhart DM. 2012. Copy number variation in the cattle genome. Funct Integr Genomics 12(4): 609–624.

Liu GE, Hou Y, Zhu B, Cardone MF, Jiang L, Cellamare A, Mitra A, Alexander LJ, Coutinho LL, Dell’Aquila ME et al. 2010. Analysis of copy number variations among diverse cattle breeds. Genome Res 20(5): 693–703.

Lorentzon M, Greenhalgh CJ, Mohan S, Alexander WS, Ohlsson C. 2005. Reduced bone mineral density in SOCS-2-deficient mice. Pediatr Res 57(2): 223–226.

Luo J, Yu Y, Mitra A, Chang S, Zhang H, Liu G, Yang N, Song J. 2013. Genome-wide copy number variant analysis in inbred chickens lines with different susceptibility to Marek’s disease. G3 (Bethesda) 3(2): 217–223.

Manolio TA, Collins FS, Cox NJ, Goldstein DB, Hindorff LA, Hunter DJ, McCarthy MI, Ramos EM, Cardon LR, Chakravarti A et al. 2009. Finding the missing heritability of complex diseases. Nature 461(7265): 747–753.

Marquez-Quinones A, Mutch DM, Debard C, Wang P, Combes M, Roussel B, Holst C, Martinez JA, Handjieva-Darlenska T, Kalouskova P et al. 2010. Adipose tissue transcriptome reflects variations between subjects with continued weight loss and subjects regaining weight 6 mo after caloric restriction independent of energy intake. Am J Clin Nutr 92(4): 975–984.

McCarroll SA, Altshuler DM. 2007. Copy-number variation and association studies of human disease. Nat Genet 39(7 Suppl): S37–42.

Metcalf D, Greenhalgh CJ, Viney E, Willson TA, Starr R, Nicola NA, Hilton DJ, Alexander WS. 2000. Gigantism in mice lacking suppressor of cytokine signalling-2. Nature 405(6790): 1069–1073.

Munoz-Amatriain M, Eichten SR, Wicker T, Richmond TA, Mascher M, Steuernagel B, Scholz U, Ariyadasa R, Spannagl M, Nussbaumer T et al. 2013. Distribution, functional impact, and origin mechanisms of copy number variation in the barley genome. Genome Biol 14(6): R58.

Murai A, Furuse M, Kitaguchi K, Kusumoto K, Nakanishi Y, Kobayashi M, Horio F. 2009. Characterization of critical factors influencing gene expression of two types of fatty acid-binding proteins (L-FABP and Lb-FABP) in the liver of birds. Comp Biochem Physiol A Mol Integr Physiol 154(2): 216–223.

Nguyen DQ, Webber C, Ponting CP. 2006. Bias of selection on human copy-number variants. PLoS Genet 2(2): e20.

Nicholas TJ, Cheng Z, Ventura M, Mealey K, Eichler EE, Akey JM. 2009. The genomic architecture of segmental duplications and associated copy number variants in dogs. Genome Res 19(3): 491–499.

Norris BJ, Whan VA. 2008. A gene duplication affecting expression of the ovine ASIP gene is responsible for white and black sheep. Genome Res 18(8): 1282–1293.

Patel RK, Jain M. 2012. NGS QC Toolkit: a toolkit for quality control of next generation sequencing data. PLoS One 7(2): e30619.

Pinto D, Darvishi K, Shi X, Rajan D, Rigler D, Fitzgerald T, Lionel AC, Thiruvahindrapuram B, Macdonald JR, Mills R et al. 2011. Comprehensive assessment of array-based platforms and calling algorithms for detection of copy number variants. Nat Biotechnol 29(6): 512–520.

Qu L, Li X, Xu G, Chen K, Yang H, Zhang L, Wu G, Hou Z, Yang N. 2006. Evaluation of genetic diversity in Chinese indigenous chicken breeds using microsatellite markers. Sci China C Life Sci 49(4): 332–341.

Redon R, Ishikawa S, Fitch KR, Feuk L, Perry GH, Andrews TD, Fiegler H, Shapero MH, Carson AR, Chen W et al. 2006. Global variation in copy number in the human genome. Nature 444(7118): 444–454.

Rosengren Pielberg G, Golovko A, Sundstrom E, Curik I, Lennartsson J, Seltenhammer MH, Druml T, Binns M, Fitzsimmons C, Lindgren G et al. 2008. A cis-acting regulatory mutation causes premature hair graying and susceptibility to melanoma in the horse. Nat Genet 40(8): 1004–1009.

Sharp AJ, Locke DP, McGrath SD, Cheng Z, Bailey JA, Vallente RU, Pertz LM, Clark RA, Schwartz S, Segraves R et al. 2005. Segmental duplications and copy-number variation in the human genome. Am J Hum Genet 77(1): 78–88.

Stranger BE, Forrest MS, Dunning M, Ingle CE, Beazley C, Thorne N, Redon R, Bird CP, de Grassi A, Lee C et al. 2007. Relative impact of nucleotide and copy number variation on gene expression phenotypes. Science 315(5813): 848–853.

Sudmant PH, Kitzman JO, Antonacci F, Alkan C, Malig M, Tsalenko A, Sampas N, Bruhn L, Shendure J, Eichler EE. 2010. Diversity of human copy number variation and multicopy genes. Science 330(6004): 641–646.

Szatkiewicz JP, Wang W, Sullivan PF, Sun W. 2013. Improving detection of copy-number variation by simultaneous bias correction and read-depth segmentation. Nucleic Acids Res 41(3):1519–1532.

Teo SM, Pawitan Y, Ku CS, Chia KS, Salim A. 2012. Statistical challenges associated with detecting copy number variations with next-generation sequencing. Bioinformatics 28(21): 2711–2718.

Tian M, Wang Y, Gu X, Feng C, Fang S, Hu X, Li N. 2013. Copy number variants in locally raised Chinese chicken genomes determined using array comparative genomic hybridization. BMC Genomics 14(1): 262.

Wang-Rodriguez J, Dreilinger AD, Alsharabi GM, Rearden A. 2002. The signaling adapter protein PINCH is up-regulated in the stroma of common cancers, notably at invasive edges. Cancer 95(6): 1387–1395.

Wang J, Jiang J, Fu W, Jiang L, Ding X, Liu JF, Zhang Q. 2012a. A genome-wide detection of copy number variations using SNP genotyping arrays in swine. BMC Genomics 13: 273.

Wang X, Nahashon S, Feaster TK, Bohannon-Stewart A, Adefope N. 2010. An initial map of chromosomal segmental copy number variations in the chicken. BMC Genomics 11: 351.

Wang Y, Gu X, Feng C, Song C, Hu X, Li N. 2012b. A genome-wide survey of copy number variation regions in various chicken breeds by array comparative genomic hybridization method. Anim Genet 43(3): 282–289.

Wapinski I, Pfeffer A, Friedman N, Regev A. 2007. Natural history and evolutionary principles of gene duplication in fungi. Nature 449(7158): 54–61.

Wong GK Liu B Wang J Zhang Y Yang X Zhang Z Meng Q Zhou J Li D Zhang J et al. 2004. A genetic variation map for chicken with 2.8 million single-nucleotide polymorphisms. Nature 432(7018): 717–722.

Wright D, Boije H, Meadows JR, Bed’hom B, Gourichon D, Vieaud A, Tixier-Boichard M, Rubin CJ, Imsland F, Hallbook F et al. 2009. Copy number variation in intron 1 of SOX5 causes the Pea-comb phenotype in chickens. PLoS Genet 5(6): e1000512.

Yalcin B, Wong K, Agam A, Goodson M, Keane TM, Gan X, Nellaker C, Goodstadt L, Nicod J, Bhomra A et al. 2011. Sequence-based characterization of structural variation in the mouse genome. Nature 477(7364): 326–329.

Yoon S, Xuan Z, Makarov V, Ye K, Sebat J. 2009. Sensitive and accurate detection of copy number variants using read depth of coverage. Genome Res 19(9): 1586–1592.

Zhang F, Gu W, Hurles ME, Lupski JR. 2009. Copy number variation in human health, disease, and evolution. Annu Rev Genomics Hum Genet 10: 451–481.

